# Paradoxical changes in brain reward status during oxycodone self-administration in a novel test of the negative reinforcement hypothesis

**DOI:** 10.1101/460048

**Authors:** Jacques D. Nguyen, Yanabel Grant, Michael A. Taffe

## Abstract

**Background and Purpose:** The extra-medical use of, and addiction to, prescription opioid analgesics is a growing health problem. To characterize how prescription opioid abuse develops, this study investigated the affective consequences of escalating prescription opioid use using intracranial self-stimulation (ICSS) reward and oxycodone intravenous self-administration (IVSA) models.

**Experimental Approach:** Male Wistar rats were given access to oxycodone IVSA (0.15 mg/kg/infusion, i.v.) in Short Access (ShA; 1 h) or Long Access (LgA; 12 h) sessions for 5 sessions/week followed by intermittent 60 h discontinuations from drug access, a novel explicit test of the negative reinforcement hypothesis. Separate groups were first trained in the ICSS procedure and then in oxycodone IVSA in 11 h LgA sessions.

**Key Results:** Rats given LgA to oxycodone escalated their responding more than ShA rats, with further significant increases observed following each 60 h discontinuation. Pre-session brain reward thresholds increased with sequential daily LgA IVSA sessions, consistent with a growing negative affective state consequent to successive daily intoxication/abstinence cycles. A 1 h oxycodone IVSA interval was sufficient to normalize these elevated reward thresholds, as was, paradoxically, a 60 h weekend abstinence. The increase in ICSS thresholds was attenuated in a group treated with the long-acting kappa opioid antagonist norBNI prior to IVSA training.

**Conclusions and Implications:** Changes in brain reward function during escalation of oxycodone self-administration are driven by an interplay between kappa opioid receptor-mediated negative affective state associated with escalated oxycodone intake and dynamic restoration of brain reward status during longer periods of abstinence.

## Introduction

Nonmedical opioid abuse is a significant global problem, with an estimated 33 million users of opiates and prescription opioids worldwide (UNODC, 2016). Approximately 2 million people in the US have a prescription opioid related abuse disorder (CBHSQ, 2015), which may increase the likelihood of later illicit opioid use (Muhuri, 2013), and prescription opioid related overdose deaths have increased by five-fold over the last two decades (CDC, 2016). Despite the growing impact of prescription opioids on public health, relatively few preclinical studies have investigated self-administration of <u>oxycodone</u>, one of the most commonly prescribed medications (OxyContin® or as part of Percocet®). The available preclinical studies have shown that mice self-administer oxycodone intravenously (Zhang, Windisch, Altschuler, Rahm, Butelman & Kreek, 2016), leading to physical dependence and withdrawal (Enga, Jackson, Damaj & Beardsley, 2016). Male and female rats express similar patterns of intake during the early stages of oxycodone self-administration training (Mavrikaki, Pravetoni, Page, Potter & Chartoff, 2017). Male rats trained to self-administer oxycodone under extended daily access conditions (8-12 h) exhibit a progressive escalation of drug intake (Matzeu & Martin-Fardon, 2020; Nguyen et al., 2019; Wade, Vendruscolo, Schlosburg, Hernandez & Koob, 2015) similar to the escalation of <u>heroin</u> self-administration observed under extended access conditions (Schlosburg et al., 2013; Vendruscolo, Schlosburg, Misra, Chen, Greenwell & Koob, 2011). Mice also exhibit escalation of oxycodone self-administration under 4 h extended access conditions (Zhang et al., 2014).

The negative reinforcement hypothesis that has been advanced to explain escalating drug IVSA under extended-access conditions (George, Koob & Vendruscolo, 2014; Koob; Koob, 2015; Koob et al., 2014; Lenoir & Ahmed, 2007) holds that the dysphoric, or negative affective, state that is experienced during the daily withdrawal of drug access grows increasingly severe with sequential sessions. The *alleviation* of this dysphoria that is provided by drug access in a subsequent session is therefore the critical stimulus which is hypothesized to increase the strength of drug-taking behavior. The intracranial self-stimulation (ICSS) reward procedure has been crucial to the argument that the behavioral escalation phenotype arises from dysregulation of common reward and affective neuronal circuitry and is not merely due to primary pharmacodynamic tolerance. For example, brain reward thresholds progressively increased with the ongoing self-administration of heroin in 23 h but not 1 h daily sessions (Kenny, Chen, Kitamura, Markou & Koob, 2006). Similarly, ICSS reward thresholds were increased compared to baseline in rats trained to self-administer <u>methamphetamine</u> in 6 h but not 1 h IVSA sessions (Jang, Whitfield, Schulteis, Koob & Wee, 2013) or <u>cocaine</u> under 6 h access (Ahmed, Kenny, Koob & Markou, 2002). Increases persisted for at least 7-8 days post-escalation, with recovery over about 10 days in the methamphetamine study. Withdrawal from <u>ethanol</u> (Schulteis, Markou, Cole & Koob, 1995), <u>nicotine</u> (Epping-Jordan, Watkins, Koob & Markou, 1998), <u>amphetamine</u> (Lin, Koob & Markou, 1999; Lin, Koob & Markou, 2000) and several opioids (Altarifi & Negus, 2011; Negus & Moerke, 2019), including <u>fentanyl</u> (Bruijnzeel, Lewis, Bajpai, Morey, Dennis & Gold, 2006), also elevates reward thresholds in rats. Correspondingly, the somatic signs of withdrawal from heroin increase progressively from 12 to 48 h in adult rats (Doherty & Frantz, 2013). These findings suggest that the motivation for drug-seeking is closely related to brain hedonic state as reflected in ICSS thresholds, and together with the behavior are predictive of abuse-related drug effects (Der-Avakian & Markou, 2012). A study by Wiebelhaus and colleagues showed that acute and repeated injections of oxycodone alter ICSS responding (Wiebelhaus, Walentiny & Beardsley, 2016); however the effects of oxycodone self-administration on ICSS brain reward thresholds have not yet been investigated. Our initial experiment for this study found that 60 h discontinuations of 12 h oxycodone access increased subsequent drug intake more than 12 h discontinuations (**Figure 1**). This frames a novel test of the negative reinforcement hypothesis through explicit manipulation of the discontinuation interval, which is a major focus of these studies.

**Figure 1.**
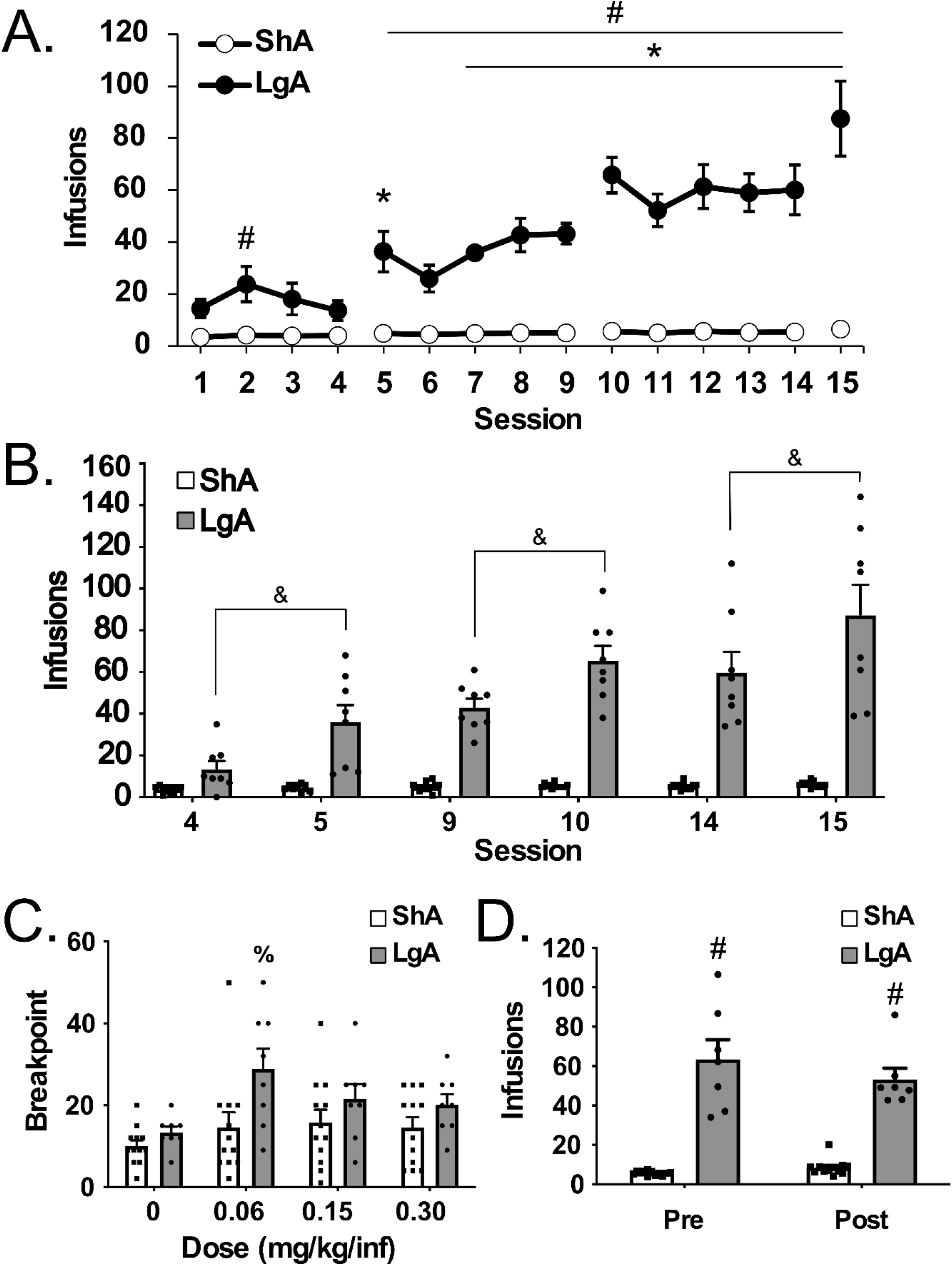
Escalation of Oxycodone Self-Administration Under Extended Access Conditions. Mean (±SEM) infusions for groups of male rats trained to self-administer of oxycodone (0.15 mg/kg/inf) within Long Access (LgA; 12 h; N=8) or Short Access (ShA; 1 h; N=12) conditions during A) acquisition (broken lines between sessions 4 & 5, 9 &10 and 14 & 15 indicate the 60 h abstinence interval); and B) before and after the weekend abstinence. C) Mean (+SEM) breakpoints reached under the progressive ratio dose-substitution procedure. D) Mean (+SEM) infusions for groups of rats before and after a 30-day abstinence period (reflects averages of the 4 sessions immediately before (Pre) and after (Post) the abstinence). Significant differences within group from session 1 are indicated by * and between group by #. Significant difference between before vs after 60 h abstinence are indicated by &. Significant difference within group from vehicle control is indicated by %.

The dynorphin / <u>kappa opioid receptor</u> (KOR) system has been shown to be one mechanism critical to the expression of anhedonia, as inferred from the ICSS procedure or from drug self-administration. For example the activation of KORs reduces the rewarding efficacy of brain stimulation (increased ICSS thresholds), whereas blockade of KORs facilitates the reversal of stress- and anhedonia-induced elevation of ICSS thresholds (Chartoff, Sawyer, Rachlin, Potter, Pliakas & Carlezon, 2012; Knoll & Carlezon, 2010; Todtenkopf, Marcus, Portoghese & Carlezon, 2004). A KOR-dependent negative affective state is critical to stress-induced potentiation of drug reward, mediating the consumption, escalation and withdrawal from drugs of abuse (Chavkin & Koob, 2016; Graziane, Polter, Briand, Pierce & Kauer, 2013; Karkhanis, Holleran & Jones, 2017; Tejeda & Bonci, 2019; Walker & Koob, 2008; Wee & Koob, 2010). More specifically, the long-lasting inactivation of KOR signaling via systemic or intracerebral administration of <u>nor-binaltorphimine</u> (norBNI) attenuates the escalation of heroin (Schlosburg et al., 2013), cocaine (Kallupi et al., 2013; Wee, Orio, Ghirmai, Cashman & Koob, 2009), alcohol (Berger, Williams, McGinnis & Walker, 2013) or methamphetamine (Whitfield et al., 2015) self-administration, under extended access conditions. These results were interpreted as a consequence of alleviating the negative affective state associated with cycles of drug-taking and (daily) discontinuation.

The present study was designed to determine if the escalation of oxycodone self-administration that is associated with extended (11-12 h) access (Wade, Vendruscolo, Schlosburg, Hernandez & Koob, 2015) is accompanied by decreased brain reward function using the ICSS procedure. The first goal was to test the hypothesis that continued cycles of extended intoxication, followed by daily abstinence intervals, facilitates an increase in drug taking driven by a progressively negative affective state. The second goal was to determine if KOR signaling contributes mechanistically to the development of this negative affective state.

## Methods and Materials

### Animals

Male Wistar rats (N=58; Charles River, Raleigh, NC) were housed in a humidity and temperature-controlled (23±1°C) vivarium on 12:12h light:dark cycles. Animals entered the laboratory at 11-14 weeks of age and weighed an average of 410.0±5.56 g at the start of the self-administration study. Animals had *ad libitum* access to food and water in their home cages and were housed in pairs throughout the study. All procedures were conducted in the animals’ scotophase, under protocols approved by the Institutional Care and Use Committees of The Scripps Research Institute and consistent with the National Institutes of Health Guide for the Care and Use of Laboratory Animals (Garber et al., 2011). These studies are reported in accordance with the ARRIVE guidelines (McGrath & Lilley, 2015). Principles of reduction, replacement and refinement were addressed in several ways in this study. Repeated-measures designs were selected for many studies to minimize the number of groups required and to enhance statistical power for comparisons. In addition, the same group was used in multiple sub-studies, thereby reducing the total number of animals required for the purpose. Subjects were randomly assigned to the major treatment groups within each experiment (Experiments 1, 2 or 3) and procedures conducted in a counter-balanced order. Treatment group sizes in Experiment 1 (N=12) and Experiment 3 (N=8) were equal in initial design and the sample size was based on power estimates based on prior literature, accounting for anticipated subject drop-out due to loss of catheter patency and lack of ICSS threshold stability during initial training/baselining. In Experiment 2, the primary goal was to assess IVSA under extended access conditions so the LgA group size (N=12) for this separate cohort was identical to that used in Experiment 1. However, variability was small in the ShA group compared to LgA in Experiment 1 and the ShA rats were used only as a negative control group, thus a smaller group (N=6) was used consistent with reduction and refinement goals. Computer automated data collection (self-administration, ICSS) were conducted in parallel across groups thus no blinding of investigator to treatment group was included in the approach. The experimental apparatus locations were balanced across the groups.

### Drugs

Oxycodone HCl was obtained from Sigma-Aldrich (St. Louis, MO) and Spectrum Chemicals (Gardena, CA). Nor-binaltorphimine dihydrochloride (norBNI), diamorphine HCL (heroin) and <u>Δ9-tetrahydrocannabinol</u> (THC) were obtained from the NIDA Drug Supply Program (Bethesda, MD). THC was supplied in ethanolic solution (50-200 mg/mL); the ethanol was evaporated off prior to preparation in a 1:1:8 ratio of ethanol:cremophor:saline vehicle (Veh). All other doses are expressed as the salt and were dissolved in physiological saline (Sal; 0.9% NaCl).

### Intravenous Catheterization and Self-Administration

Procedures were adapted from protocols previously described (Nguyen et al., 2019; Nguyen, Grant, Creehan, Vandewater & Taffe, 2017; Nguyen, Hwang, Grant, Janda & Taffe, 2018), and a detailed summary of the experimental design can be found in the **Supplementary Materials**.

Rats (Experiment 1) were implanted with chronic indwelling catheters under anesthesia with an <u>isoflurane</u>/oxygen vapor mixture (isoflurane 5% induction, 1-3% maintenance) and were then trained daily (Monday-Friday) to self-administer intravenous oxycodone (0.15 mg/kg/infusion; ∼0.1 ml/infusion) under Long Access (LgA; 12 h) or Short Access (ShA; 1 h) conditions with 60 h intervals of discontinuation across weekends. The dose was chosen from precedent literature demonstrating robust acquisition and behavioral escalation under extended access conditions (Nguyen et al., 2019; Wade, Vendruscolo, Schlosburg, Hernandez & Koob, 2015). Following acquisition, rats were subjected to randomized Progressive-Ratio (PR) dose-response testing wherein doses of oxycodone (0-0.3 mg/kg/infusion) were presented in a balanced order on sequential sessions lasting up to 3 h. Rats were returned to their home cages for an extended 30-day abstinence period and then returned to 12 h sessions to test for re-engagement of drug seeking (re-escalation) following detoxification.

### Intracranial Self-Stimulation (ICSS) Reward

For these studies, procedures were adapted from well-established protocols describe (Kenny & Markou, 2006; Kornetsky & Esposito, 1979; Markou & Koob, 1992; Nguyen, Aarde, Cole, Vandewater, Grant & Taffe, 2016). Rats were anesthetized with an isoflurane/oxygen vapor mixture (isoflurane 5% induction, 1-3% maintenance) and prepared with unilateral electrodes aimed at the medial forebrain bundle (coordinates: AP −0.5mm, ML ±1.7mm, DV skull −9.5mm). Rats were trained in a procedure adapted from the discrete-trial current-threshold procedure (Kenny & Markou, 2006; Kornetsky & Esposito, 1979; Markou & Koob, 1992; Nguyen, Aarde, Cole, Vandewater, Grant & Taffe, 2016) until stable ICSS thresholds were exhibited. Thereafter, the rats were anesthetized with an isoflurane/oxygen vapor mixture (isoflurane 5% induction, 1-3% maintenance), surgically implanted with intravenous catheters, allowed to recover for a minimum of 1 week. ICSS training was resumed for at least one week to re-establish baseline thresholds and thereafter rats were randomly assigned to ShA (1 h) or to LgA (11 h) self-administration conditions. Rats in Experiment 2 completed daily (M-F) self-administration sessions after ICSS sessions for sequential weeks with weekend discontinuations (60 h). In Experiment 3, a separate group of rats was trained in the ICSS procedure and injected with norBNI (30 mg/kg, i.p.; LgA-norBNI) or the saline vehicle (LgA-sal) 3 days prior to the beginning of oxycodone self-administration training. The norBNI dose was chosen from precedent literature demonstrating robust effects on behavioral escalation under extended access conditions for heroin, cocaine or methamphetamine self-administration across intervals of time similar to the acquisition interval of the present experiment (Schlosburg et al., 2013; Wee, Orio, Ghirmai, Cashman & Koob, 2009; Whitfield et al., 2015). Effects of norBNI are predicted to last about 21-28 days in mice (Bruchas et al., 2007; Melief et al., 2011) and rats (Jones & Holtzman, 1992). In Experiment 3 a two week abstinence period was imposed between sessions 15 and 16. Injections of THC were administered 30 min prior to the start of self-administration. Additional experimental details can be found in the **Supplementary Materials**.

### Data Analysis

Analysis of the IVSA data was conducted with Analysis of Variance (ANOVA) on the number of infusions earned (ShA: 1 h session; LgA: 11 or 12 h session), on the breakpoints reached in the PR study and on the percent change in brain reward threshold (µA) relative to individual baseline. ICSS data were normalized to individual baselines due to wide individual variation in raw µA values common to this procedure. An animal (N=1 rat in Experiment 3) was excluded from ICSS analysis because it failed to exhibit a decreased reward threshold after a non-contingent methamphetamine injection (0.5 mg/kg, i.p). This was used as a positive control based on the robustness of this response in prior investigations (Nguyen, Aarde, Cole, Vandewater, Grant & Taffe, 2016). Within-subjects factors of Session, Drug Dose (PR) and/or pretreatment condition were included as warranted and between-subjects factors for Access Duration and saline/norBNI pretreatment. Post hoc analysis, using Tukey (multi-level factors) or Sidak (two-level factors) tests to control for multiple comparisons, followed only after any significant main effects or interactions from the ANOVA. Group sizes for a given analysis cell refer to independent rat observations. Data analysis was not undertaken for any group sizes less than N<5. All statistics were performed in Prism (Graphpad Software, Inc, La Jolla, CA). In all cases, p<0.05 was the criterion for statistical significance.

## Results

### Escalation of oxycodone self-administration under extended access conditions

In Experiment 1, mean number of intravenous oxycodone infusions obtained by rats trained under LgA (N=8) conditions was significantly higher than infusions obtained by ShA (N=12) rats (**Figure1A**). More infusions were obtained by LgA rats during sessions 5 and 7-15 compared to session 1, and LgA rats received significantly more infusions compared to ShA rats during sessions 5-15 and that drug intake did not change across sessions for the ShA group. Analysis of the first 10, 30 and 60 minutes of self-administration confirmed escalation in the LgA group but not in the ShA group (see **Supplementary Figure S1**). During the final session of acquisition, LgA rats exhibited 76.9% drug-associated lever responding. The ShA group exhibited >80% drug-associated lever responding for 9 of the final 11 days of acquisition. Analysis of the sessions before and after the 60 h weekend abstinence periods showed that LgA rats significantly increased drug intake following this extended drug deprivation, whereas ShA showed no significant change (**Figure 1B**). The LgA and ShA rats also exhibited group differences during PR dose substitution (**Figure 1C**). Neither LgA (N=7) nor ShA (N=12) rats exhibited any significant change in oxycodone infusions (0.15 mg/kg/inf) obtained under a FR1 response contingency (**Figure 1D**) before versus after the 30-day abstinence interval. One rat (LgA) that completed the acquisition interval was not included in the post-abstinence assessment due to illness. Significantly more infusions were obtained by LgA rats on the day after a weekend break.

### Brain stimulation reward during escalation of oxycodone self-administration under extended access conditions

The LgA and ShA rats in the Experiment 2 (ICSS-trained) cohort also exhibited group differences in oxycodone self-administration (**Figure 2A**). More infusions were obtained by the LgA group in sessions 14, 15, 19-24, 26-28 compared with the first session. In addition, significant group differences were confirmed across weekend breaks (Sessions 11-17 and 19-28) (**Figure 2B**). For LgA rats the analysis of the weekend effects confirmed significant increases for each Monday session over the prior end of week session from the 11^th^ session to the 24^th^. Whereas for ShA rats, mean infusions were not significantly different between Pre-weekend and Monday sessions (**Figure 2C**). Three rats were excluded due to opioid related self-injury during acquisition (N=1) or catheter failure, thus LgA (N=10) and ShA (N=5).

**Figure 2.**
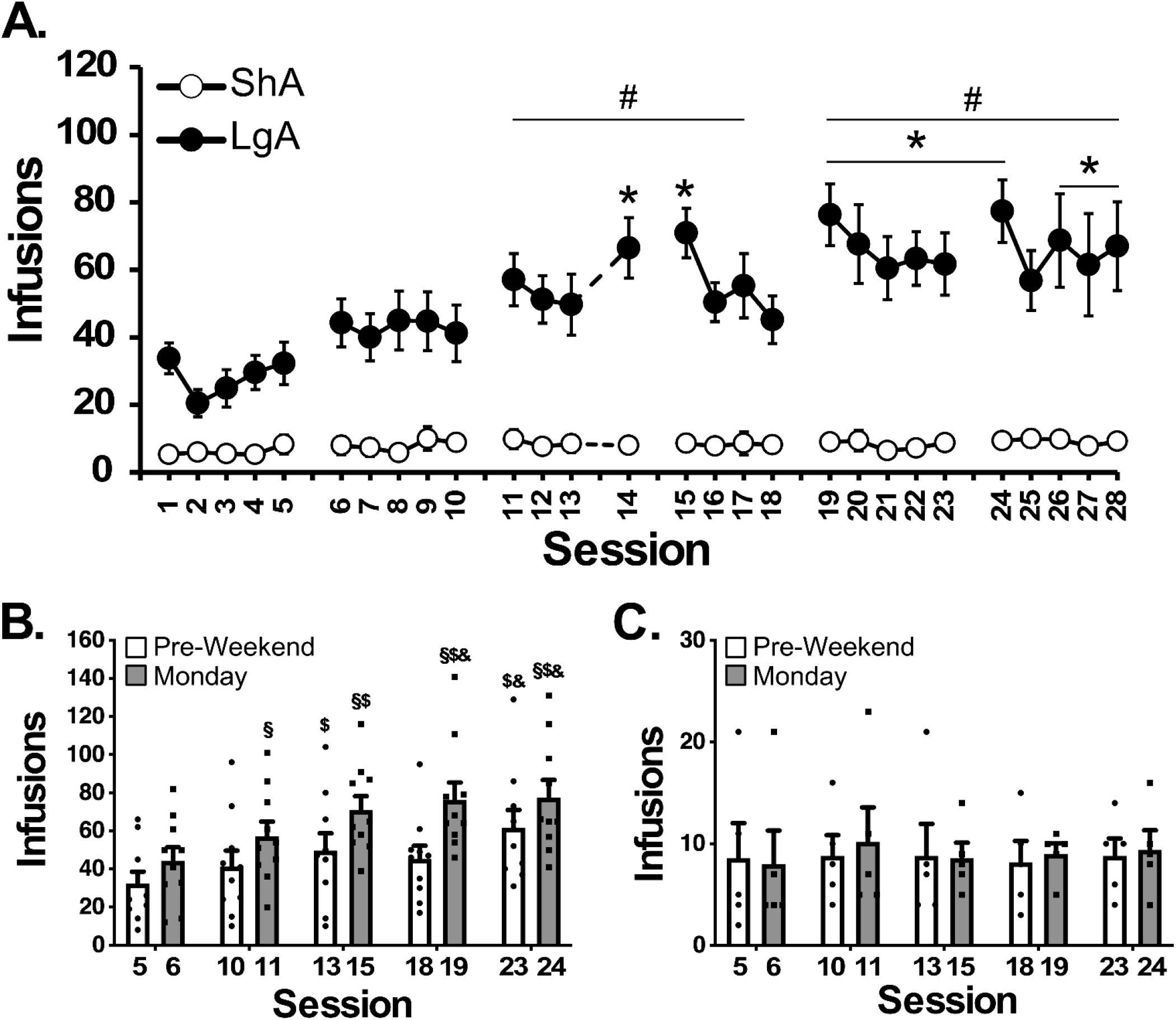
Escalation of Oxycodone Self-Administration in ICSS-trained rats. A) Mean (±SEM) infusions for groups of rats trained to self-administer of oxycodone (0.15 mg/kg/inf) in Long Access (LgA; 11 h; N=10) or Short Access (ShA; 1 h; N=5) sessions following the daily ICSS evaluation. A significant difference from the first session is indicated with * and a significant difference compared with the corresponding day of the ShA group by #. Breaks in the series indicate the weekend and the dotted line indicates a Thursday on which the self-administration session was omitted. Mean (±SEM) infusions for LgA (B) and ShA (C) across weekend breaks. A significant difference across the weekend is indicated with § and significant differences within week from the first week by $ and from the second week by &.

ICSS thresholds in the LgA group in Experiment 2 increased across successive days in each week of self-administration but returned nearly to baseline across the weekend breaks (**Figure 3A**). Analysis confirmed significant increases within each week of self-administration, no change within the abstinence week and differences between days of the self-administration weeks compared with the corresponding weekday of the abstinence week. An attenuation of the ICSS threshold elevation was observed on the 5^th^ day of the third week, following omission of the IVSA session the previous day (see **Figure 2A**).

**Figure 3.**
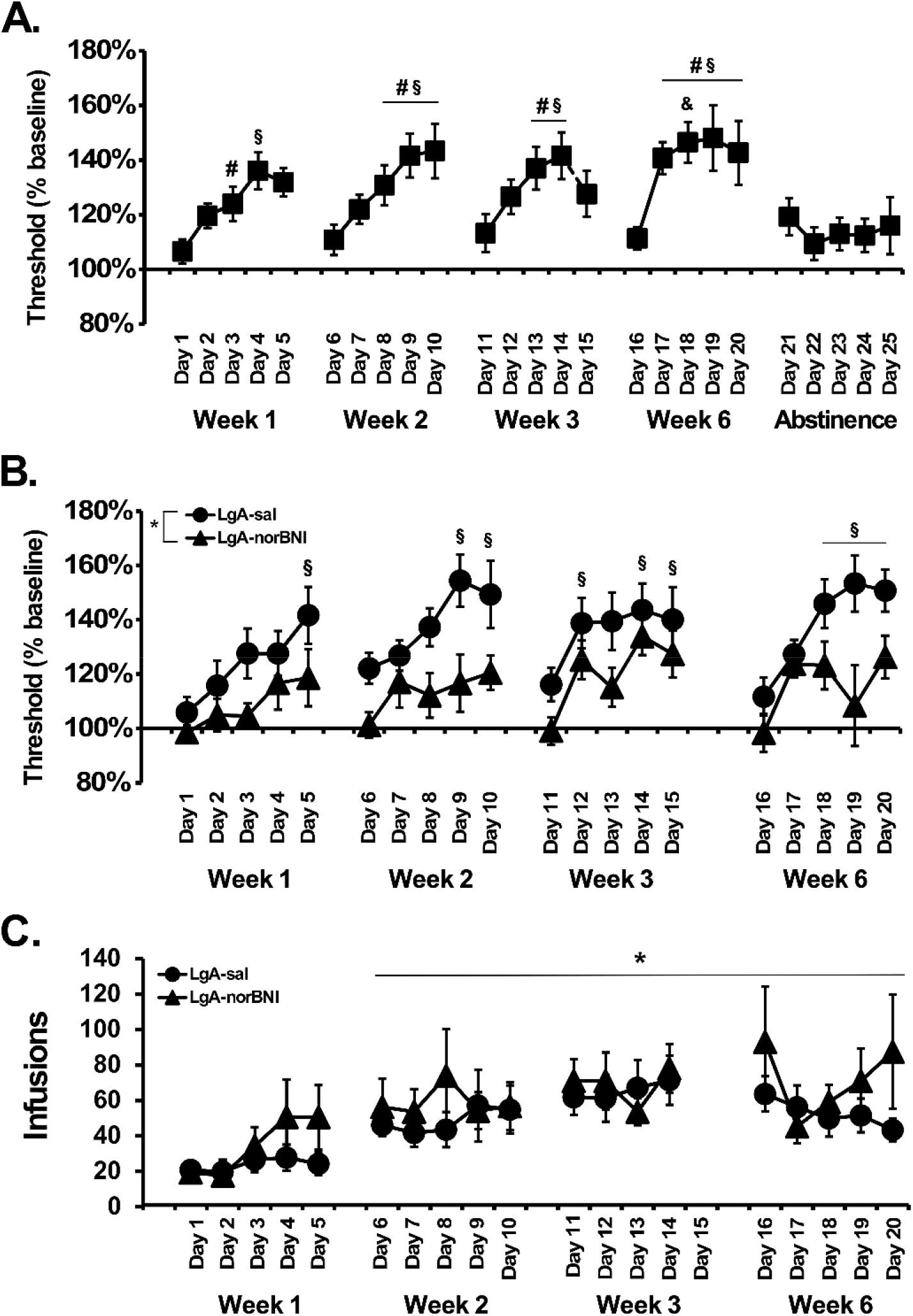
Oxycodone self-administration alters brain reward thresholds partially via kappa opioid receptor signaling. A) Mean (N=10; ±SEM) ICSS thresholds expressed as percent of the individual baseline threshold in long access rats. A significant difference from the first day of the week is indicated by §, and a significant difference compared with the corresponding day of the Abstinence week by # and from Week 1 by &. Analysis excluded the week 3, day 15 session before which IVSA was omitted. B) Mean ICSS thresholds in LgA-sal (N=8; ±SEM) and LgA-norBNI (N=7; ±SEM) rats. A significant difference between group is indicated by *. A significant difference, collapsed across groups, from the first day of that week is indicated by §. C) Mean (±SEM) oxycodone infusions in LgA (N=8) and LgA-norBNI (N=7) rats. A significant difference, collapsed across groups, from Day 1 is indicated by *.

In Experiment 3, groups of rats (LgA-norBNI) that were injected with norBNI (30 mg/kg, i.p.), a kappa opioid receptor (KOR) antagonist, or saline (LgA-sal), prior to initiating self-administration training were used to determine the effect of KOR function on oxycodone-induced effects on ICSS reward thresholds (**Figure 3B**). Analysis confirmed that compared to LgA-sal control rats, LgA-norBNI rats had significantly lower ICSS thresholds across successive days (Monday through Friday) during 4 weeks of self-administration. ICSS reward thresholds, collapsed across groups, were increased (Days 5, 9-10, 12, 14-15, and 18-20) compared to the first day of the respective week. Interestingly, mean oxycodone infusions did not differ between LgA-sal control rats or LgA-norBNI rats (**Figure 3C**). Session 15 was excluded from this analysis, as the self-administration session was only 60 minutes followed by an ICSS session. Analysis confirmed that infusions, collapsed across groups, were significantly increased (Day 6-20) compared to the first day of IVSA training.

We further confirmed in Experiment 3 that the within-week elevations of brain reward thresholds depended on access duration and generalized across oxycodone and heroin (**Figure 4A**). ICSS thresholds were elevated following seven sequential IVSA sessions (Days 16-23) in both LgA-sal and LgA-norBNI rats but returned to the baseline following four consecutive 11 h sessions of saline self-administration (Days 24-27). Subsequent oxycodone IVSA in 1 h access sessions in Week 6 (Days 28-31) failed to significantly increase reward thresholds. Heroin (0.06 mg/kg/infusion) IVSA under 11 h access duration (Days 32-41) significantly elevated reward thresholds in both groups in Week 7 and then in 4 h access sessions in Week 8. There were significant changes relative to pre-saline (Day 23) and the first day of each heroin week, respectively. Group comparisons of daily ICSS thresholds showed consistency across the LgA (Experiment 2) and LgA-saline (Experiment 3) groups, as they exhibited very similar elevations of reward thresholds across the initial three weeks (**Figure 4B**). More interestingly, the pattern of ICSS threshold increases in the LgA-norBNI (Experiment 3) group was very similar to the pattern of modest increases observed in the ShA (Experiment 2) group. Analysis comparing the ICSS threshold changes within the four groups across the first 14 sessions confirmed a main effect of Group and of Session; however, the Dunnett post hoc test of the marginal means did not confirm any significant difference between LgA and any of the other three groups.

**Figure 4.**
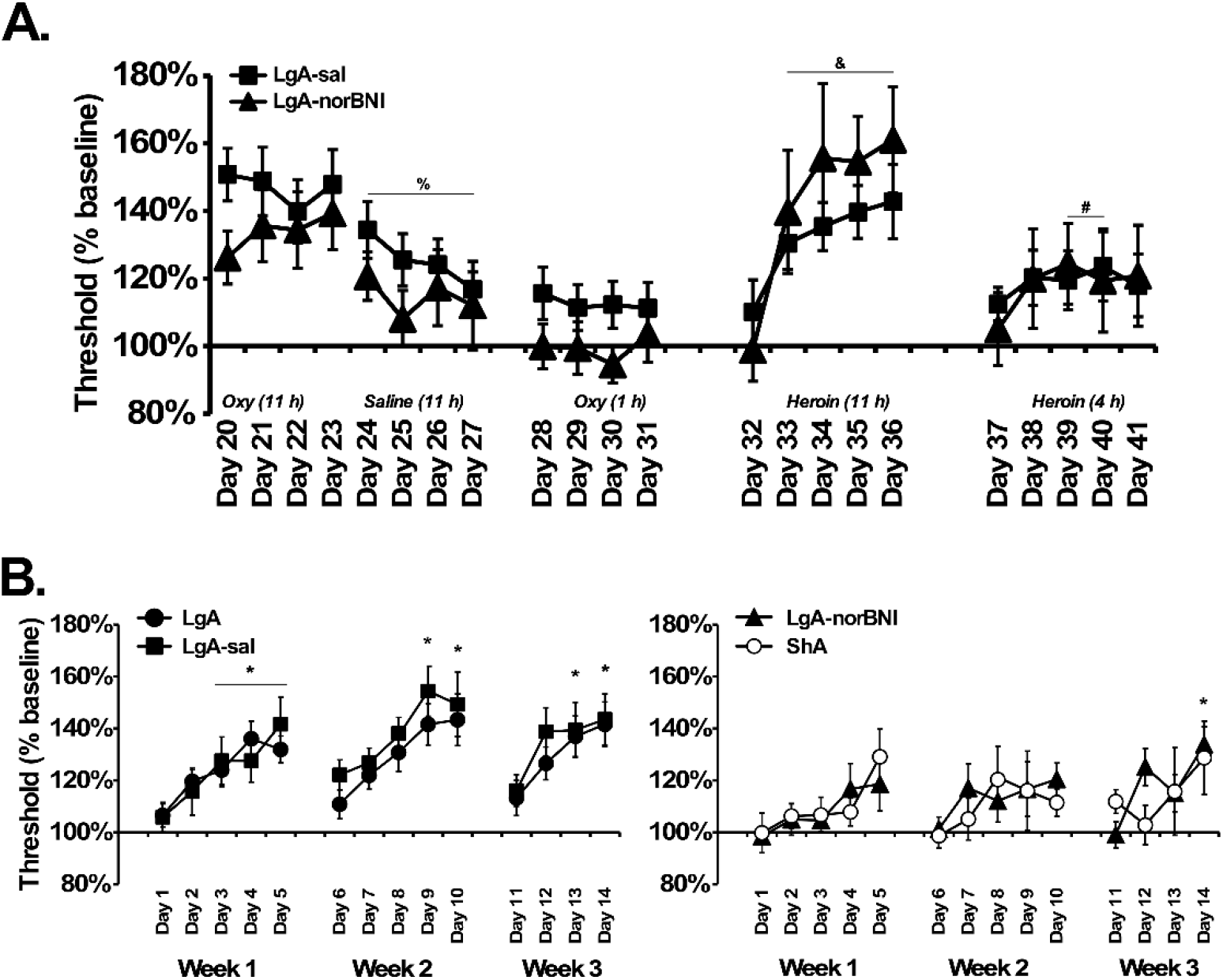
Elevation in brain reward threshold is differentially affected by drug identity and access duration. A) Mean ICSS thresholds in LgA-sal (N=8; ±SEM) and LgA-norBNI (N=7; ±SEM) rats expressed as percent of the individual baseline threshold. A significant difference, collapsed across group, from Day 23 is indicated with %, from Day 32 with & and from Day 37 with #. B) Group comparisons of threshold elevations during oxycodone self-administration confirmed that the Long Access group from Experiment 2 (LgA; 12 h) and the Long Access group from Experiment 3 (LgA-sal; 11 h; saline pretreatment) group exhibited similar and robust ICSS threshold elevations across the first three weeks of IVSA, and that the Short Access (ShA; 1 h) and Long Access group pretreated with nor-BNI (LgA-norBNI; 11 h) exhibited similarly limited ICSS threshold elevations across the first three weeks of IVSA. A significant difference from the first day of the week, across groups, is indicated with *.

### Brain stimulation reward procedure does not affect oxycodone self-administration

To determine if the ICSS procedure prior to the IVSA sessions impacted oxycodone intake, the acquisition data for the four LgA cohorts was compared using the days of access that were shared across all four groups. This meant that the fourth day of the third week was omitted (because it was omitted for the animals in Experiment 2), the fifth day of the third week was omitted (because it was only a one hour session for the Experiment 3 groups) and one day of the first week was omitted because Experiment 1 started on a four day week. This analysis confirmed a significant effect of Session but did not confirm any Group differences or interaction. During Sessions 51-54 for the Experiment 3 groups, the ICSS session was omitted twice, and included twice, in counter balanced order to determine any effects on oxycodone self-administration. No significant effect of omitting the ICSS session was confirmed (see **Supplementary Figure S4**).

### Normalization of elevated ICSS thresholds by a one hour self-administration session

Reward thresholds were assessed before and after a 1 h IVSA session in LgA groups in Experiments 2 (**Figure 5A**) and 3 (**Figure 5B-D**). In the Experiment 2 group, the analysis of the ICSS thresholds before and after a 1 h oxycodone IVSA session on Monday and Friday confirmed that pre-IVSA reward thresholds were significantly higher on Friday compared with Monday and that a significant reduction in threshold was observed after an hour of self-administration (**Figure 5A**). Analysis of ICSS thresholds in Experiment 3 also confirmed a significant reduction after a 1 h self-administration session (**Figure 5B**) with no difference in this effect between LgA and LgA-norBNI groups. In contrast, there was no significant threshold reduction caused by a 1 h self-administration session when the groups were limited to 1 h sessions for the entire week (**Figure 5C**). Finally, there was a significant threshold reduction after 1 h of heroin IVSA during the daily 11 h heroin IVSA training (**Figure 5D**), again with no difference between LgA and LgA-norBNI groups.

**Figure 5.**
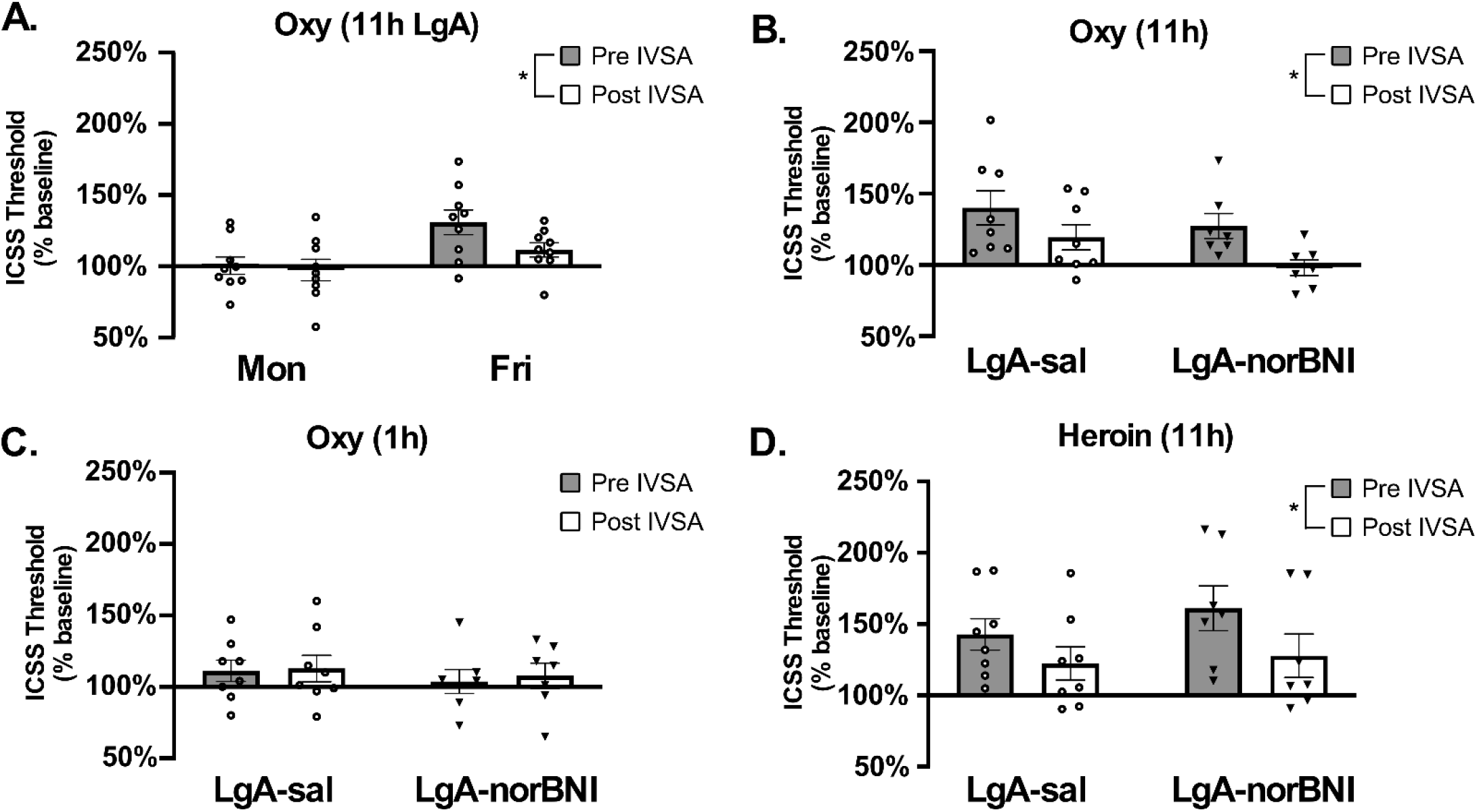
Normalization of ICSS thresholds by 1 h IVSA of oxycodone or heroin. A) Mean (N=9; ±SEM) ICSS thresholds before and after IVSA sessions. Mean (N=7-8; ±SEM) thresholds before and after B,C) 1 h IVSA of oxycodone or D) 1 h IVSA of heroin in rats trained under 1 (C) and/or 11 h (A, B, D) conditions. A significant marginal mean difference is indicated by *.

### Combined reinforcer efficacy versus overall oxycodone intake

In this study rats were injected daily with THC (5 mg/kg, i.p.) or vehicle prior to 6 h IVSA sessions for four sequential days; pretreatments were counterbalanced across two weeks within the total sample (i.e., LgA-sal and LgA-norBNI combined). Our previous study showed that THC decreases oxycodone self-administration (Nguyen et al., 2019) by enhancing the rewarding efficacy of a given unit dose. Data from the current experiment confirmed that this reduction in oxycodone self-administration behavior caused by THC (**Figure 6A**) did not change the pattern of within-week increase in ICSS thresholds (**Figure 6B**). Analysis of the self-administration data confirmed that infusions were significantly different between rats injected with vehicle (Veh) or THC (5 mg/kg, i.p.) across days (**Figure 6A**). Analysis of ICSS data confirmed that reward thresholds significantly increased in the rats when treated with either vehicle or THC (**Figure 6B**). As the 30-minute bin analysis of the self-administration pattern shows (**Figure 6C**), the THC condition blunted, but did not eliminate, intake during the loading phase and the first several hours.

**Figure 6.**
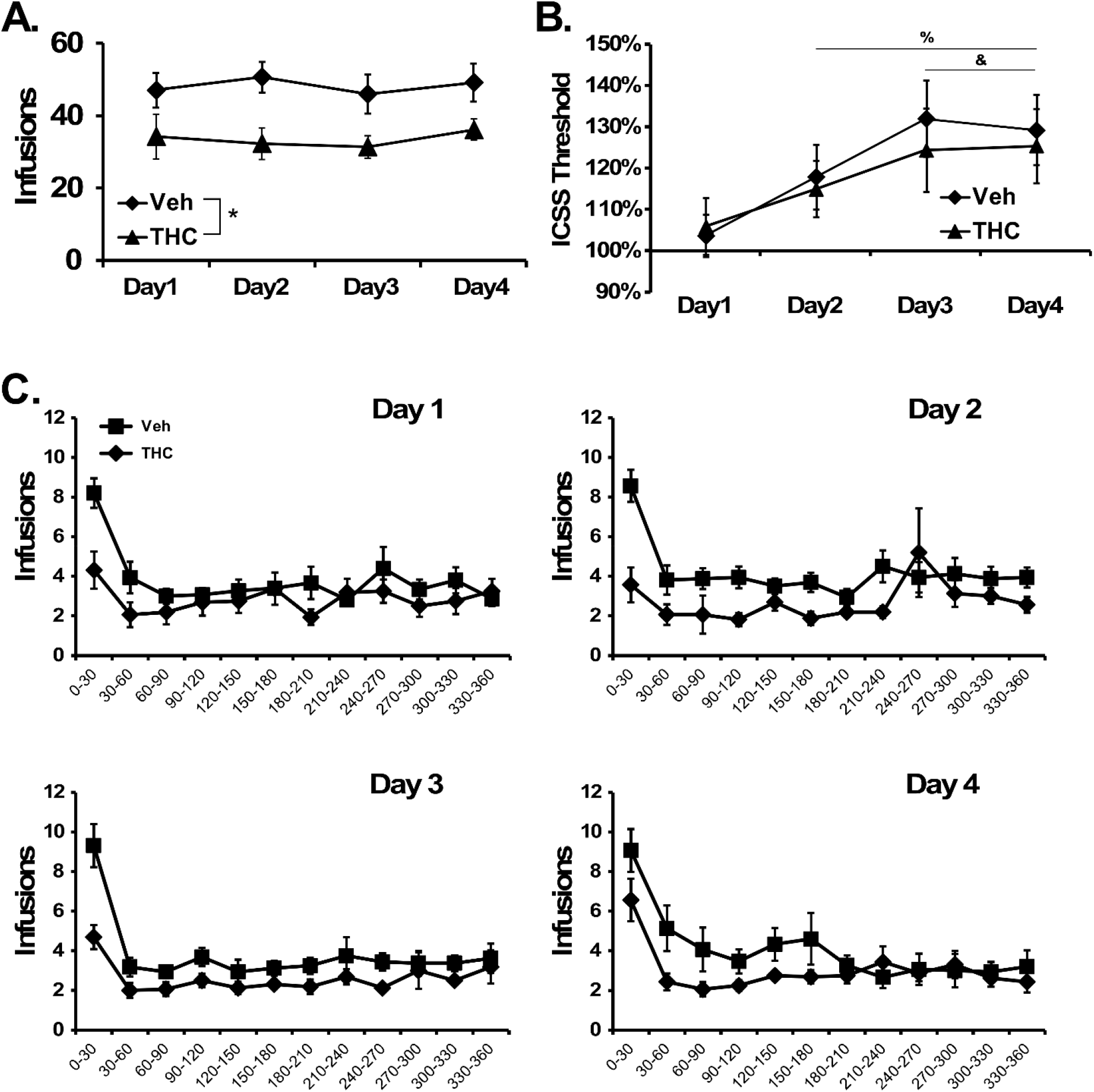
Effect of THC on oxycodone self-administration and subsequent ICSS threshold determinations. A) Mean (±SEM) oxycodone infusions obtained by all (N=16) rats under 6 h extended access conditions with vehicle or THC (5 mg/kg, i.p.) injected prior to the session for four consecutive days, in a balanced order, on two successive weeks. A significant difference between pre-treatment conditions is indicated with *. B) Mean (±SEM) ICSS thresholds by rats (N=15) determined prior to injection on each day. A significant difference compared with Day 1, across treatments, is indicated with % and compared with Day 2, across treatments, with &. C) Mean oxycodone infusions obtained on each day, represented as 30-minute bins across the session, for each pre-treatment condition.

## Discussion

Diversion of prescription opioid medications such as oxycodone for nonmedical use, and the transition to abuse of illicit opioids has had a tremendous negative impact on public health and safety worldwide over the past two decades. Intravenous self-administration (IVSA) procedures using laboratory rodents have recently been used to model oxycodone-seeking behavior (Blackwood, McCoy, Ladenheim & Cadet, 2019; Ginsburg & Lamb, 2018; Matzeu & Martin-Fardon, 2020; Mavrikaki, Pravetoni, Page, Potter & Chartoff, 2017), to elucidate the effects of oxycodone abuse on cellular mechanisms and synaptic function (Blackwood et al., 2019; Yuferov, Zhang, Liang, Zhao, Randesi & Kreek, 2018; Zhang et al., 2017; Zhang, Liang, Randesi, Yuferov, Zhao & Kreek, 2018), and to develop and assess potential therapeutic interventions for oxycodone abuse (Neelakantan et al., 2017; Nguyen, Hwang, Grant, Janda & Taffe, 2018; Townsend et al., 2017). Prior investigations showed that long (12 h or 3 consecutive 3 h) daily access to oxycodone in IVSA procedures resulted in progressively escalated drug-taking compared with shorter (1 h) access (Blackwood et al., 2019; Wade, Vendruscolo, Schlosburg, Hernandez & Koob, 2015). In this study, we confirm those findings since rats that were provided access to oxycodone in 12 h sessions increased their mean intake across weeks of training, whereas animals allowed 1 h access sessions did not. The dose-response pattern in the PR procedure further confirmed the elevated efficacy of oxycodone as a reinforcer in Long-Access (LgA) versus Short-Access (ShA) rats, without a change in potency. (This latter finding is consistent with prior investigations of extended-access drug self-administration and is inconsistent with explanations based on pharmacological tolerance where a rightward shift of the dose-effect function would be expected (Zernig et al., 2007)). This qualitative outcome is consistent with that reported by Wade and colleagues; any quantitative differences in the breakpoints achieved and the dose of greatest LgA/ShA differentiation are likely due to minor procedural differences such as the PR schedules employed and the between-versus within-group approach to the dose-effect manipulation. Importantly, we discovered (as first reported in pre-print form; (Nguyen et al., 2017)) that intermittent 60 h deprivations further increased oxycodone IVSA, producing a step-wise, upward ratcheting pattern of abstinence-related drug seeking (**Figure 1**).

The behavioral data support an interpretation that escalated oxycodone IVSA under long access conditions is driven in part by increasing negative affect associated with longer durations of drug abstinence. This represents a novel test of the negative reinforcement hypothesis. Although so called “deprivation effects” have been previously reported for the self-administration of other drugs of abuse under some conditions, such studies have not explicitly manipulated the deprivation interval to determine if the rate of escalation is affected by increasing one or more of the inter-session interval(s). For example, Heyser and colleagues showed that ethanol intake increases after a period of 3-28 days of deprivation compared to baseline (Heyser, Schulteis & Koob, 1997). Intermittent 24-48 h of abstinence produced an escalation of nicotine intake (Cohen, Koob & George, 2012), particularly in the first deprivation period. Intermittent access followed by 7 day abstinence during the early stages of cocaine IVSA significantly potentiated drug seeking (Calipari, Siciliano, Zimmer & Jones, 2015). Since the investigation was only conducted in males, the findings reported in this paper cannot be assumed to generalize to female rats. Sex differences in oral (Fulenwider et al., 2020; Zanni, DeSalle, Deutsch, Barr & Eisch, 2020) and intravenous (Kimbrough et al., 2020; Nguyen, Creehan, Kerr & Taffe, 2020) oxycodone intake have been reported in rats, although another study found no sex difference in sustained IVSA of oxycodone under varying training conditions (Mavrikaki, Pravetoni, Page, Potter & Chartoff, 2017). Interestingly, even when sex differences in IVSA under fixed-ratio conditions were reported, there were no sex differences under a progressive ratio dose substitution (Nguyen, Creehan, Kerr & Taffe, 2020) or in withdrawal-induced behaviors (Kimbrough et al., 2020), suggesting similar motivation and similar induction of negative affect across sexes. Finally, Mavrikaki and colleagues have recently shown that prior to exposure to chronic morphine injections may differentially affect acquisition of oxycodone IVSA in males compared to females (Mavrikaki, Lintz, Constantino, Page & Chartoff, 2020).

The increased ICSS thresholds observed during sequential daily extended-access self-administration sessions in this study were consistent with prior results showing brain reward changes following extended access to self-administration of several different drugs (Ahmed, Kenny, Koob & Markou, 2002; Jang, Whitfield, Schulteis, Koob & Wee, 2013; Kenny, Chen, Kitamura, Markou & Koob, 2006); this similarity is emphasized in the eight consecutive day experiment (**Figure 4A**). This pattern has been interpreted as key support for the negative reinforcement hypothesis in those reports, also see (Pantazis, Gonzalez, Tunstall, Carmack, Koob & Vendruscolo, 2021; Zernig et al., 2007). The interpretation that dependence-related negative affect drove the escalated oxycodone IVSA in *this* study is further supported by the finding that when ICSS thresholds were elevated, the self-administration of oxycodone (or heroin) for one hour (**Figure 5**), or non-contingent injection of an appropriate dose of oxycodone (**Figure S2**), was able to normalize thresholds towards baseline. As in prior studies, the elevations in reward thresholds produced by long access to oxycodone were more robust than those produced by short access IVSA, as evidenced by both the ShA group (**Figure 4B**) and the 1 h access experiments in the LgA groups (**Figure 4A**). Similarly, elevations were greater during 11 h heroin access versus 4 h access (**Figure 4B**) and during 11 h oxycodone access compared with 6 h access (**Figure 3A vs Figure 6B**). Reward threshold elevations may also depend on the rewarding efficacy of a given drug, in addition to the specific access duration. For example, self-administration of the highly efficacious reinforcer (Aarde, Huang, Creehan, Dickerson & Taffe, 2013; Collins, Sulima, Rice & France, 2019; Watterson et al., 2014) 3,4-methylenedioxypyrovalerone (MDPV) in 1 h sessions elevated reward thresholds across 10 days (Geste, Pompilus, Febo & Bruijnzeel, 2018). In a related vein, the reinforcing efficacy of a unit dose of oxycodone self-administered by rats is enhanced by THC (Nguyen et al., 2019) and we similarly found in this study that THC pretreatment reduced oxycodone intake in a 6 h IVSA session (**Figure 6A**) by attenuating, but not eliminating, the loading dose phase (**Figure 6C**). The current study showed that reduced oxycodone intake, produced by THC pretreatment, did not alter the increase in reward threshold that was observed across 4 sequential IVSA sessions (**Figure 6B**). Interestingly, these data complement the effect of norBNI in Experiment 3 in which a difference in the ICSS pattern was observed without any difference in oxycodone self-administration, and further illustrate the partial independence of these indices. The THC effect is likely due to an increase in the reinforcing efficacy of a unit dose of oxycodone, as in our prior study (Nguyen et al., 2019). The rats do exhibit a loading phase and continue to respond for drug infusions in a regular pattern. Thus, THC is not merely suppressing behavior or, likely, the drug seeking. The opioid-sparing effects of THC are further extended by our finding that repeated THC exposure during adolescence may alter motivation for fentanyl reward during adulthood (Nguyen, Creehan, Kerr & Taffe, 2020).

Correlational analyses of the relationship between oxycodone infusions and ICSS reward thresholds following extended abstinence (i.e. between Sessions 5-6, 10-11, 13-15, 18-19) in this study failed to confirm a significant Pearson correlation (**Figure Supplemental Figure S5**). Analysis of the cumulative number of oxycodone infusions each week failed to confirm a correlation with ICSS thresholds (**Figure Supplemental Figure S6**). Furthermore, analysis of relative changes in ICSS thresholds across the week failed to confirm a correlation with self-administration (**Figure Supplemental Figure S7**). As a caveat, the number of oxycodone infusions across the last 2 days of week 1 showed a significant correlation with reduction in ICSS reward threshold. These analyses suggest that differences in drug intake may not predict alterations in ICSS reward threshold nor the converse on an individual basis, particularly with modest sample sizes predicated on group effects. Overall, a lack of strong correlation between ICSS reward thresholds and oxycodone infusions is consistent with the “paradoxical” observation based on established interpretative value of these models.

The most interesting (i.e., paradoxical) feature of the present findings was the re-setting of ICSS thresholds to baseline levels, combined with the upward ratcheting of oxycodone IVSA, after 60 h of abstinence. The trend established by 24 h of discontinuation (**Figure 3A**; Day 15) suggested a linear recovery of brain reward over several days, which was later confirmed by the saline substitution experiment (**Figure 4A**). This latter finding was consistent with the ∼week-long recovery of ICSS thresholds observed following cessation of methamphetamine LgA IVSA (Jang, Whitfield, Schulteis, Koob & Wee, 2013). Relatedly, a period of 23.5 h abstinence following chronic morphine injections reduced ICSS reward (Altarifi & Negus, 2011) and somatic and behavioral signs of withdrawal were reported 12 h after the last LgA oxycodone (0.15 mg/kg/infusion, 12 h) self-administration session in recent studies (de Guglielmo, Kallupi, Sedighim, Newman & George, 2020; Kallupi et al., 2020; Kimbrough et al., 2020). The failure of 1 h of oxycodone IVSA to alter brain reward on the first session after 60 h discontinuation (**Figure 5A**, also see **Figure S2**) is an important clue. This supports the conclusion that as dependence progresses, drug-taking is no longer effective as a euphoriant but is sufficient only to *normalize* affect. It implies that self-administration is governed by a *change* in brain reward from an allostatic set-point (Koob, 2015), rather than a fixed target relative to a basal threshold set prior to the onset of drug experience. (As a minor caveat, traditional measures of physical dependence were not obtained in this study, but see above regarding recent findings using similar self-administration protocols). Recovery of brain reward systems is possible, since reward thresholds were significantly decreased below baseline by non-contingent injection of oxycodone, or the non-opioid drug methamphetamine, (see **Figure S3**) after a long interval of discontinuation (3-4 weeks). Similarly, food-maintained responding that was decreased by chronic morphine and further inhibited by drug discontinuation, returned to the levels observed during chronic morphine administration over about 3 days, but required two weeks to fully return to pre-morphine baselines (Ford & Balster, 1976).

Prior work shows that activation of kappa opioid receptor (KOR) signaling elevates ICSS thresholds and decreases motivation to respond for brain stimulation reward (Conway, Puttick, Russell, Potter, Roitman & Chartoff, 2019; Faunce & Banks, 2020; Russell et al., 2014; Todtenkopf, Marcus, Portoghese & Carlezon, 2004). Negus and colleagues demonstrated a lack of effect of the kappa antagonist norBNI on *pain-induced depression* of ICSS (Negus, Morrissey, Rosenberg, Cheng & Rice, 2010); this contrasts with the effect of norBNI pretreatment in the present data where it attenuated *oxycodone-withdrawal-induced elevations* of ICSS brain reward thresholds. It has also been shown that norBNI can attenuate an increase in ICSS thresholds that is caused by withdrawal from daily noncontingent injections of cocaine (Chartoff, Sawyer, Rachlin, Potter, Pliakas & Carlezon, 2012). Interestingly, <u>desipramine</u> normalized ICSS threshold elevations induced by cocaine withdrawal (Markou, Hauger & Koob, 1992), an effect that may be related to the ability of this tricyclic antidepressant to reduce stress-related dynorphin expression (Chartoff et al., 2009) or to directly activate KORs (Onali, Dedoni & Olianas, 2010).

In the present study, brain reward threshold elevations observed in rats treated with systemic norBNI (30 mg/kg, i.p.; LgA-norBNI) before the start of LgA IVSA were significantly reduced compared to LgA-sal rats (**Figure 3B**) and paralleled the modest and inconsistent elevations observed in ShA rats (**Figure 4**). The effects of norBNI appeared to last about six weeks, extending the previous 21-28 day interval reported for other behavioral effects in rats (Jones & Holtzman, 1992) and mice (Bruchas et al., 2007; Melief et al., 2011). Interestingly, ICSS thresholds in LgA-sal rats were higher at the beginning of each week compared to LgA-norBNI rats, suggesting only a *partial* normalization of reward function after 60 h of discontinuation in the LgA-sal rats. Work that shows lasting efficacy of norBNI in rats, for a variety of behavioral endpoints, has generally required doses of 20-30 mg/kg, i.p., (Harshberger, Gilson, Gillett, Stone, El Amrani & Valdez, 2016; Jarman, Haney & Valdez, 2018). There may be a threshold as low as 10 mg/kg, i.p. or s.c., for efficacy on some measures in rats (Anderson, Agoglia, Morales, Varlinskaya & Spear, 2013; Grella, Funk, Coen, Li & Lê, 2014; Karkhanis, Rose, Weiner & Jones, 2016; Wille-Bille, Ferreyra, Sciangula, Chiner, Nizhnikov & Pautassi, 2017). While Walker and colleagues used 5 mg/kg, s.c. per injection in rats, it required 3 daily doses to observe a decrease in alcohol self-administration (Walker, Zorrilla & Koob, 2011). In a separate study, norBNI was administered at a dose of 5 mg/kg, i.p. twice at a 12 h interval, and it was unclear that this dose is sufficient to cause comparable effects (Hölter, Henniger, Lipkowski & Spanagel, 2000). Finally, Morales et al. 2014 used doses of norBNI from 2.5 to 10 mg/kg and showed only 10 mg/kg had efficacy in rats (Morales, Anderson, Spear & Varlinskaya, 2014). Overall, the data from our study confirm that the increases in ICSS brain reward thresholds caused by sequential days of LgA IVSA of oxycodone are mediated in part by kappa opioid receptor signaling.

The present study suggests that a more nuanced view of the relationship between drug discontinuation, motivation for drug-taking and affective state *as indexed by brain reward threshold* is needed. Brain reward thresholds were normalized after 60 hours of discontinuation, while self-administration was increased. This might be viewed as paradoxical, or inconsistent with the negative affect hypothesis as an explanation of escalated drug self-administration. However, the brain reward thresholds were not *changed* by oxycodone self-administration after 60 h discontinuation. This insensitivity to change of brain reward status in the euphoric direction might explain increased self-administration behavior. Similarly, the relatively stable levels of self-administration throughout the week might reflect the fact that when thresholds were elevated after only 12 h withdrawal, one hour of oxycodone self-administration was sufficient to acutely reduce reward threshold. This outcome suggests that it is the relative change in reward threshold subsequent to drug taking that is the more important regulator of self-administration compared with brain reward status at the start of a given self-administration session.

## Conclusions

In summary, this study suggests that escalation of oxycodone self-administration under extended access with intermittent longer abstinence periods is mediated in part by negative reinforcement processes in a time-dependent manner. Overall, we conclude that drug access and discontinuation intervals each impact the acquisition and maintenance of the self-administration of oxycodone. Importantly, these findings may have clinically relevant implications. For example, these data would suggest that individuals on short-term prescribed oxycodone regimens do not skip prescribed days of treatment in an attempt to ‘tough it out’, as this may in fact lead to increased liability for abuse. These findings highlight the importance of adherence monitoring or adherence enhancing interventions, as non-adherence to pain medication use is very common (Timmerman, Stronks, Groeneweg & Huygen, 2016b) and particularly because there does not appear to be an association between medication adherence and pain treatment outcome (Timmerman, Stronks, Groeneweg & Huygen, 2016a). These data further suggest that a lack of medication adherence may increase a liability for the early stages of oxycodone addiction.

## Author Contributions

J.D.N. and M.A.T. designed the studies. J.D.N. and Y.G. performed the research and conducted initial data analysis. J.D.N. and M.A.T. conducted the statistical analysis of data, created figures, and wrote the paper. All authors approved of submitted version of the manuscript.

## Acknowledgments

This work was funded by support from the United States Public Health Service National Institutes of Health grants R01 DA035281 (MAT) and K99 DA047413 (JDN). The National Institutes of Health / NIDA had no direct influence on the design, conduct, analysis or decision to publication of the findings.

## Competing Financial Interests

The authors declare no competing financial interests that influenced the conduct or reporting of this work.

### Abbreviations

ICSS: intracranial self-stimulation
IVSA: intravenous self-administration
LgA: Long Access
ShA: Short Access
PR: progressive ratio
norBNI: nor-binaltorphimine
KOR: kappa opioid receptor

## References

Aarde SM, Huang PK, Creehan KM, Dickerson TJ, & Taffe MA (2013). The novel recreational drug 3,4-methylenedioxypyrovalerone (MDPV) is a potent psychomotor stimulant: self-administration and locomotor activity in rats. Neuropharmacology 71: 130–140.

Ahmed SH, Kenny PJ, Koob GF, & Markou A (2002). Neurobiological evidence for hedonic allostasis associated with escalating cocaine use. Nat Neurosci 5: 625–626.

Altarifi AA, & Negus SS (2011). Some determinants of morphine effects on intracranial self-stimulation in rats: dose, pretreatment time, repeated treatment, and rate dependence. Behav Pharmacol 22: 663–673.

Anderson RI, Agoglia AE, Morales M, Varlinskaya EI, & Spear LP (2013). Stress, κ manipulations, and aversive effects of ethanol in adolescent and adult male rats. Neuroscience 249: 214–222.

Berger AL, Williams AM, McGinnis MM, & Walker BM (2013). Affective cue-induced escalation of alcohol self-administration and increased 22-kHz ultrasonic vocalizations during alcohol withdrawal: role of kappa-opioid receptors. Neuropsychopharmacology 38: 647–654.

Blackwood CA, Hoerle R, Leary M, Schroeder J, Job MO, McCoy MT, et al. (2019). Molecular Adaptations in the Rat Dorsal Striatum and Hippocampus Following Abstinence-Induced Incubation of Drug Seeking After Escalated Oxycodone Self-Administration. Mol Neurobiol 56: 3603–3615.

Blackwood CA, McCoy MT, Ladenheim B, & Cadet JL (2019). Escalated Oxycodone Self-Administration and Punishment: Differential Expression of Opioid Receptors and Immediate Early Genes in the Rat Dorsal Striatum and Prefrontal Cortex. Front Neurosci 13: 1392.

Bruchas MR, Yang T, Schreiber S, Defino M, Kwan SC, Li S, et al. (2007). Long-acting kappa opioid antagonists disrupt receptor signaling and produce noncompetitive effects by activating c-Jun N-terminal kinase. J Biol Chem 282: 29803–29811.

Bruijnzeel AW, Lewis B, Bajpai LK, Morey TE, Dennis DM, & Gold M (2006). Severe deficit in brain reward function associated with fentanyl withdrawal in rats. Biol Psychiatry 59: 477–480.

Calipari ES, Siciliano CA, Zimmer BA, & Jones SR (2015). Brief intermittent cocaine self-administration and abstinence sensitizes cocaine effects on the dopamine transporter and increases drug seeking. Neuropsychopharmacology 40: 728–735.

CBHSQ (2015). 2014 National Survey on Drug Use and Health: Detailed Tablesed. Administration S.A.a.M.H.S.: Rockville, MD.

Wide-ranging online data for epidemiologic research (WONDER). [Online] Available from http://wonder.cdc.gov. [Accessed: 2019].

Chartoff E, Sawyer A, Rachlin A, Potter D, Pliakas A, & Carlezon WA (2012). Blockade of kappa opioid receptors attenuates the development of depressive-like behaviors induced by cocaine withdrawal in rats. Neuropharmacology 62: 167–176.

Chartoff EH, Papadopoulou M, MacDonald ML, Parsegian A, Potter D, Konradi C, et al. (2009). Desipramine reduces stress-activated dynorphin expression and CREB phosphorylation in NAc tissue. Mol Pharmacol 75: 704–712.

Chavkin C, & Koob GF (2016). Dynorphin, Dysphoria, and Dependence: the Stress of Addiction. Neuropsychopharmacology 41: 373–374.

Cohen A, Koob GF, & George O (2012). Robust escalation of nicotine intake with extended access to nicotine self-administration and intermittent periods of abstinence. Neuropsychopharmacology 37: 2153–2160.

Collins GT, Sulima A, Rice KC, & France CP (2019). Self-administration of the synthetic cathinones 3,4-methylenedioxypyrovalerone (MDPV) and alpha-pyrrolidinopentiophenone (alpha-PVP) in rhesus monkeys. Psychopharmacology (Berl) 236: 3677–3685.

Conway SM, Puttick D, Russell S, Potter D, Roitman MF, & Chartoff EH (2019). Females are less sensitive than males to the motivational-and dopamine-suppressing effects of kappa opioid receptor activation. Neuropharmacology 146: 231–241.

de Guglielmo G, Kallupi M, Sedighim S, Newman AH, & George O (2020). Dopamine D3 Receptor Antagonism Reverses the Escalation of Oxycodone Self-administration and Decreases Withdrawal-Induced Hyperalgesia and Irritability-Like Behavior in Oxycodone-Dependent Heterogeneous Stock Rats. Frontiers in behavioral neuroscience 13.

Der-Avakian A, & Markou A (2012). The neurobiology of anhedonia and other reward-related deficits. Trends in neurosciences 35: 68–77.

Doherty JM, & Frantz KJ (2013). Attenuated effects of experimenter-administered heroin in adolescent vs. adult male rats: physical withdrawal and locomotor sensitization. Psychopharmacology (Berl) 225: 595–604.

Enga RM, Jackson A, Damaj MI, & Beardsley PM (2016). Oxycodone physical dependence and its oral self-administration in C57BL/6J mice. Eur J Pharmacol 789: 75–80.

Epping-Jordan MP, Watkins SS, Koob GF, & Markou A (1998). Dramatic decreases in brain reward function during nicotine withdrawal. Nature 393: 76–79.

Faunce KE, & Banks ML (2020). Effects of repeated kappa-opioid receptor agonist U-50488 treatment and subsequent termination on intracranial self-stimulation in male and female rats. Exp Clin Psychopharmacol 28: 44–54.

Ford RD, & Balster RL (1976). Schedule-controlled behavior in the morphine-dependent rat. Pharmacol Biochem Behav 4: 569–573.

Fulenwider HD, Nennig SE, Hafeez H, Price ME, Baruffaldi F, Pravetoni M, et al. (2020). Sex differences in oral oxycodone self-administration and stress-primed reinstatement in rats. Addict Biol 25: e12822.

Garber JC, Barbee RW, Bielitzki JT, Clayton LA, Donovan JC, Hendriksen CFM, et al. (2011) Guide for the Care and Use of Laboratory Animals, 8th Edition. National Academies Press: Washington D.C.

George O, Koob GF, & Vendruscolo LF (2014). Negative reinforcement via motivational withdrawal is the driving force behind the transition to addiction. Psychopharmacology (Berl) 231: 3911–3917.

Geste JR, Pompilus M, Febo M, & Bruijnzeel AW (2018). Self-administration of the synthetic cathinone MDPV enhances reward function via a nicotinic receptor dependent mechanism. Neuropharmacology 137: 286–296.

Ginsburg BC, & Lamb RJ (2018). Frustration stress (unexpected loss of alternative reinforcement) increases opioid self-administration in a model of recovery. Drug Alcohol Depend 182: 33–39.

Graziane NM, Polter AM, Briand LA, Pierce RC, & Kauer JA (2013). Kappa opioid receptors regulate stress-induced cocaine seeking and synaptic plasticity. Neuron 77: 942–954.

Grella SL, Funk D, Coen K, Li Z, & Lê Ad (2014). Role of the kappa-opioid receptor system in stress-induced reinstatement of nicotine seeking in rats. Behav Brain Res 265: 188–197.

Harshberger E, Gilson EA, Gillett K, Stone JH, El Amrani L, & Valdez GR (2016). nor-BNI Antagonism of Kappa Opioid Agonist-Induced Reinstatement of Ethanol-Seeking Behavior. J Addict 2016: 1084235.

Heyser CJ, Schulteis G, & Koob GF (1997). Increased ethanol self-administration after a period of imposed ethanol deprivation in rats trained in a limited access paradigm. Alcohol Clin Exp Res 21: 784–791.

Hölter SM, Henniger MS, Lipkowski AW, & Spanagel R (2000). Kappa-opioid receptors and relapse-like drinking in long-term ethanol-experienced rats. Psychopharmacology (Berl) 153: 93–102.

Jang CG, Whitfield T, Schulteis G, Koob GF, & Wee S (2013). A dysphoric-like state during early withdrawal from extended access to methamphetamine self-administration in rats. Psychopharmacology (Berl) 225: 753–763.

Jarman SK, Haney AM, & Valdez GR (2018). Kappa opioid regulation of depressive-like behavior during acute withdrawal and protracted abstinence from ethanol. PLoS One 13: e0205016.

Jones DN, & Holtzman SG (1992). Long term kappa-opioid receptor blockade following nor-binaltorphimine. Eur J Pharmacol 215: 345–348.

Kallupi M, Carrette LLG, Kononoff J, Solberg Woods LC, Palmer AA, Schweitzer P, et al. (2020). Nociceptin attenuates the escalation of oxycodone self-administration by normalizing CeA-GABA transmission in highly addicted rats. Proc Natl Acad Sci U S A.

Kallupi M, Wee S, Edwards S, Whitfield TW, Jr., Oleata CS, Luu G, et al. (2013). Kappa opioid receptor-mediated dysregulation of gamma-aminobutyric acidergic transmission in the central amygdala in cocaine addiction. Biol Psychiatry 74: 520–528.

Karkhanis A, Holleran KM, & Jones SR (2017). Dynorphin/Kappa Opioid Receptor Signaling in Preclinical Models of Alcohol, Drug, and Food Addiction. Int Rev Neurobiol 136: 53–88.

Karkhanis AN, Rose JH, Weiner JL, & Jones SR (2016). Early-Life Social Isolation Stress Increases Kappa Opioid Receptor Responsiveness and Downregulates the Dopamine System. Neuropsychopharmacology 41: 2263–2274.

Kenny PJ, Chen SA, Kitamura O, Markou A, & Koob GF (2006). Conditioned withdrawal drives heroin consumption and decreases reward sensitivity. J Neurosci 26: 5894–5900.

Kenny PJ, & Markou A (2006). Nicotine self-administration acutely activates brain reward systems and induces a long-lasting increase in reward sensitivity. Neuropsychopharmacology 31: 1203–1211.

Kimbrough A, Kononoff J, Simpson S, Kallupi M, Sedighim S, Palomino K, et al. (2020). Oxycodone self-administration and withdrawal behaviors in male and female Wistar rats. Psychopharmacology (Berl) 237: 1545–1555.

Knoll AT, & Carlezon WA, Jr. (2010). Dynorphin, stress, and depression. Brain Res 1314: 56–73.

Koob GF Neurobiology of Opioid Addiction: Opponent Process, Hyperkatifeia and Negative Reinforcement. Biological Psychiatry.

Koob GF (2015). The dark side of emotion: the addiction perspective. Eur J Pharmacol 753: 73–87.

Koob GF, Buck CL, Cohen A, Edwards S, Park PE, Schlosburg JE, et al. (2014). Addiction as a stress surfeit disorder. Neuropharmacology 76 Pt B: 370–382.

Kornetsky C, & Esposito RU (1979). Euphorigenic drugs: effects on the reward pathways of the brain. Federation proceedings 38: 2473–2476.

Lenoir M, & Ahmed SH (2007). Heroin-induced reinstatement is specific to compulsive heroin use and dissociable from heroin reward and sensitization. Neuropsychopharmacology 32: 616–624.

Lin D, Koob GF, & Markou A (1999). Differential effects of withdrawal from chronic amphetamine or fluoxetine administration on brain stimulation reward in the rat--interactions between the two drugs. Psychopharmacology (Berl) 145: 283–294.

Lin D, Koob GF, & Markou A (2000). Time-dependent alterations in ICSS thresholds associated with repeated amphetamine administrations. Pharmacol Biochem Behav 65: 407–417.

Markou A, Hauger RL, & Koob GF (1992). Desmethylimipramine attenuates cocaine withdrawal in rats. Psychopharmacology (Berl) 109: 305–314.

Markou A, & Koob GF (1992). Construct validity of a self-stimulation threshold paradigm: effects of reward and performance manipulations. Physiology & behavior 51: 111–119.

Matzeu A, & Martin-Fardon R (2020). Targeting the orexin system for prescription opioid use disorder: Orexin-1 receptor blockade prevents oxycodone taking and seeking in rats. Neuropharmacology 164: 107906.

Mavrikaki M, Lintz T, Constantino N, Page S, & Chartoff E (2020). Chronic opioid exposure differentially modulates oxycodone self-administration in male and female rats. Addict Biol: e12973.

Mavrikaki M, Pravetoni M, Page S, Potter D, & Chartoff E (2017). Oxycodone self-administration in male and female rats. Psychopharmacology (Berl) 234: 977–987.

McGrath JC, & Lilley E (2015). Implementing guidelines on reporting research using animals (ARRIVE etc.): new requirements for publication in BJP. Br J Pharmacol 172: 3189–3193.

Melief EJ, Miyatake M, Carroll FI, Beguin C, Carlezon WA, Jr., Cohen BM, et al. (2011). Duration of action of a broad range of selective kappa-opioid receptor antagonists is positively correlated with c-Jun N-terminal kinase-1 activation. Mol Pharmacol 80: 920–929.

Morales M, Anderson RI, Spear LP, & Varlinskaya EI (2014). Effects of the kappa opioid receptor antagonist, nor-binaltorphimine, on ethanol intake: impact of age and sex. Dev Psychobiol 56: 700–712.

Muhuri PKG, J. C.; Davies, M. C. (2013). Associations of Nonmedical Pain Reliever Use and Initiation of Heroin Use in the United States.

Neelakantan H, Holliday ED, Fox RG, Stutz SJ, Comer SD, Haney M, et al. (2017). Lorcaserin Suppresses Oxycodone Self-Administration and Relapse Vulnerability in Rats. ACS chemical neuroscience 8: 1065–1073.

Negus SS, & Moerke MJ (2019). Determinants of opioid abuse potential: Insights using intracranial self-stimulation. Peptides 112: 23–31.

Negus SS, Morrissey EM, Rosenberg M, Cheng K, & Rice KC (2010). Effects of kappa opioids in an assay of pain-depressed intracranial self-stimulation in rats. Psychopharmacology (Berl) 210: 149–159.

Nguyen JD, Aarde SM, Cole M, Vandewater SA, Grant Y, & Taffe MA (2016). Locomotor Stimulant and Rewarding Effects of Inhaling Methamphetamine, MDPV, and Mephedrone via Electronic Cigarette-Type Technology. Neuropsychopharmacology 41: 2759–2771.

Nguyen JD, Creehan KM, Kerr TM, & Taffe MA (2020). Lasting effects of repeated Δ(9)-tetrahydrocannabinol vapour inhalation during adolescence in male and female rats. Br J Pharmacol 177: 188–203.

Nguyen JD, Grant Y, Creehan KM, Hwang CS, Vandewater SA, Janda KD, et al. (2019). Delta(9)-tetrahydrocannabinol attenuates oxycodone self-administration under extended access conditions. Neuropharmacology 151: 127–135.

Nguyen JD, Grant Y, Creehan KM, Vandewater SA, & Taffe MA (2017). Escalation of intravenous self-administration of methylone and mephedrone under extended access conditions. Addiction biology 22: 1160–1168.

Nguyen JD, Hwang CS, Grant Y, Janda KD, & Taffe MA (2018). Prophylactic vaccination protects against the development of oxycodone self-administration. Neuropharmacology 138: 292–303.

Nguyen JD, Kirson D, Steinman MQ, Patel R, Khom S, Varodayan FP, et al. (2017). Withdrawal-induced escalated oxycodone self-administration is mediated by kappa opioid receptor function. bioRxiv.

Onali P, Dedoni S, & Olianas MC (2010). Direct agonist activity of tricyclic antidepressants at distinct opioid receptor subtypes. J Pharmacol Exp Ther 332: 255–265.

Pantazis CB, Gonzalez LA, Tunstall BJ, Carmack SA, Koob GF, & Vendruscolo LF (2021). Cues conditioned to withdrawal and negative reinforcement: Neglected but key motivational elements driving opioid addiction. Science Advances 7: eabf0364.

Russell SE, Rachlin AB, Smith KL, Muschamp J, Berry L, Zhao Z, et al. (2014). Sex differences in sensitivity to the depressive-like effects of the kappa opioid receptor agonist U-50488 in rats. Biol Psychiatry 76: 213–222.

Schlosburg JE, Whitfield TW, Jr., Park PE, Crawford EF, George O, Vendruscolo LF, et al. (2013). Long-term antagonism of kappa opioid receptors prevents escalation of and increased motivation for heroin intake. J Neurosci 33: 19384–19392.

Schulteis G, Markou A, Cole M, & Koob GF (1995). Decreased brain reward produced by ethanol withdrawal. Proc Natl Acad Sci U S A 92: 5880–5884.

Tejeda HA, & Bonci A (2019). Dynorphin/kappa-opioid receptor control of dopamine dynamics: Implications for negative affective states and psychiatric disorders. Brain Res 1713: 91–101.

Timmerman L, Stronks DL, Groeneweg G, & Huygen FJ (2016a). The Value of Medication-Specific Education on Medication Adherence and Treatment Outcome in Patients with Chronic Pain: A Randomized Clinical Trial. Pain medicine (Malden, Mass) 17: 1829–1837.

Timmerman L, Stronks DL, Groeneweg JG, & Huygen FJ (2016b). Prevalence and determinants of medication non-adherence in chronic pain patients: a systematic review. Acta anaesthesiologica Scandinavica 60: 416–431.

Todtenkopf MS, Marcus JF, Portoghese PS, & Carlezon WA, Jr. (2004). Effects of kappa-opioid receptor ligands on intracranial self-stimulation in rats. Psychopharmacology (Berl) 172: 463–470.

Townsend EA, Naylor JE, Negus SS, Edwards SR, Qureshi HN, McLendon HW, et al. (2017). Effects of nalfurafine on the reinforcing, thermal antinociceptive, and respiratory-depressant effects of oxycodone: modeling an abuse-deterrent opioid analgesic in rats. Psychopharmacology (Berl) 234: 2597–2605.

UNODC (2016). World Drug ReportUnited Nations Office on Drugs and Crime.

Vendruscolo LF, Schlosburg JE, Misra KK, Chen SA, Greenwell TN, & Koob GF (2011). Escalation patterns of varying periods of heroin access. Pharmacol Biochem Behav 98: 570–574.

Wade CL, Vendruscolo LF, Schlosburg JE, Hernandez DO, & Koob GF (2015). Compulsive-like responding for opioid analgesics in rats with extended access. Neuropsychopharmacology 40: 421–428.

Walker BM, & Koob GF (2008). Pharmacological evidence for a motivational role of kappa-opioid systems in ethanol dependence. Neuropsychopharmacology 33: 643–652.

Walker BM, Zorrilla EP, & Koob GF (2011). Systemic kappa-opioid receptor antagonism by nor-binaltorphimine reduces dependence-induced excessive alcohol self-administration in rats. Addict Biol 16: 116–119.

Watterson LR, Kufahl PR, Nemirovsky NE, Sewalia K, Grabenauer M, Thomas BF, et al. (2014). Potent rewarding and reinforcing effects of the synthetic cathinone 3,4-methylenedioxypyrovalerone (MDPV). Addict Biol 19: 165–174.

Wee S, & Koob GF (2010). The role of the dynorphin-kappa opioid system in the reinforcing effects of drugs of abuse. Psychopharmacology (Berl) 210: 121–135.

Wee S, Orio L, Ghirmai S, Cashman JR, & Koob GF (2009). Inhibition of kappa opioid receptors attenuated increased cocaine intake in rats with extended access to cocaine. Psychopharmacology (Berl) 205: 565–575.

Whitfield TW, Jr., Schlosburg JE, Wee S, Gould A, George O, Grant Y, et al. (2015). kappa Opioid receptors in the nucleus accumbens shell mediate escalation of methamphetamine intake. J Neurosci 35: 4296–4305.

Wiebelhaus JM, Walentiny DM, & Beardsley PM (2016). Effects of Acute and Repeated Administration of Oxycodone and Naloxone-Precipitated Withdrawal on Intracranial Self-Stimulation in Rats. J Pharmacol Exp Ther 356: 43–52.

Wille-Bille A, Ferreyra A, Sciangula M, Chiner F, Nizhnikov ME, & Pautassi RM (2017). Restraint stress enhances alcohol intake in adolescent female rats but reduces alcohol intake in adolescent male and adult female rats. Behav Brain Res 332: 269–279.

Yuferov V, Zhang Y, Liang Y, Zhao C, Randesi M, & Kreek MJ (2018). Oxycodone Self-Administration Induces Alterations in Expression of Integrin, Semaphorin and Ephrin Genes in the Mouse Striatum. Front Psychiatry 9: 257.

Zanni G, DeSalle MJ, Deutsch HM, Barr GA, & Eisch AJ (2020). Female and male rats readily consume and prefer oxycodone to water in a chronic, continuous access, two-bottle oral voluntary paradigm. Neuropharmacology 167: 107978.

Zernig G, Ahmed SH, Cardinal RN, Morgan D, Acquas E, Foltin RW, et al. (2007). Explaining the escalation of drug use in substance dependence: models and appropriate animal laboratory tests. Pharmacology 80: 65–119.

Zhang Y, Liang Y, Levran O, Randesi M, Yuferov V, Zhao C, et al. (2017). Alterations of expression of inflammation/immune-related genes in the dorsal and ventral striatum of adult C57BL/6J mice following chronic oxycodone self-administration: a RNA sequencing study. Psychopharmacology (Berl) 234: 2259–2275.

Zhang Y, Liang Y, Randesi M, Yuferov V, Zhao C, & Kreek MJ (2018). Chronic Oxycodone Self-administration Altered Reward-related Genes in the Ventral and Dorsal Striatum of C57BL/6J Mice: An RNA-seq Analysis. Neuroscience 393: 333–349.

Zhang Y, Mayer-Blackwell B, Schlussman SD, Randesi M, Butelman ER, Ho A, et al. (2014). Extended access oxycodone self-administration and neurotransmitter receptor gene expression in the dorsal striatum of adult C57BL/6 J mice. Psychopharmacology (Berl) 231: 1277–1287.

Zhang Y, Windisch K, Altschuler J, Rahm S, Butelman ER, & Kreek MJ (2016). Adolescent oxycodone self administration alters subsequent oxycodone-induced conditioned place preference and anti-nociceptive effect in C57BL/6J mice in adulthood. Neuropharmacology 111: 314–322.

