## Supplemental Materials for "Paradoxical changes in brain reward status during oxycodone self-administration in a novel test of the negative reinforcement hypothesis"

***Supplementary Information***

### Methods and Materials

**Drugs.** Oxycodone HCl and d-methamphetamine HCl were obtained from Sigma-Aldrich (St. Louis, MO) and Spectrum Chemicals (Gardena, CA). Nor-binaltorphimine dihydrochloride (norBNI) was obtained from the NIDA Drug Supply Program (Bethesda, MD). Doses are expressed as the salt and were dissolved in physiological saline (0.9% NaCl). Acute oxycodone and methamphetamine injections were administered 15 min prior to the start of ICSS, and THC injections were administered 30 min prior to the start of self-administration.

**Intravenous Catheterization.** Rats were anesthetized with an isoflurane/oxygen vapor mixture (isoflurane 5% induction, 1-3% maintenance) and prepared with chronic indwelling intravenous catheters as described previously (Nguyen, Grant, Creehan, Vandewater & Taffe, 2017). Briefly, the intravenous catheters consisted of a 14.5-cm length of polyurethane based tubing (Micro-Renathane®, Braintree Scientific, Inc, Braintree, MA) fitted to a guide cannula (Plastics One, Roanoke, VA) curved at an angle and encased in dental cement anchored to an ~3 cm circle of durable mesh. Catheter tubing was passed subcutaneously from the animal's back to the right jugular vein. Catheter tubing was inserted into the vein and tied gently with suture thread. A liquid tissue adhesive was used to close the incisions (3M™ Vetbond™ Tissue Adhesive: 1469SB, 3M, St. Paul, MN). A minimum of 4 days was allowed for surgical recovery prior to starting an experiment. For the first three days of the recovery period, an antibiotic (cefazolin) and an analgesic (flunixin) were administered daily. During testing and training, intravenous catheters were flushed with ~0.2-0.3 ml heparinized (166.7 USP/ml) saline before sessions and ~0.2-0.3 ml heparinized saline containing cefazolin (100 mg/mL) after sessions. Catheter patency was assessed once a week after the last session of the week, via administration through the catheter of ~0.2 ml (10 mg/ml) of the ultra-short-acting barbiturate anesthetic Brevital sodium (1% methohexital sodium; Eli Lilly, Indianapolis, IN). Animals with patent catheters exhibit prominent signs of anesthesia (pronounced loss of muscle tone) within 3 sec after infusion. Animals that failed to display these signs were considered to have faulty catheters, and if catheter patency failure was detected, data that were collected after the previous passing of this test were excluded from analysis.

**Self-Administration Procedures.** Drug self-administration was conducted in operant boxes (Med Associates, Inc., Fairfax, VT) located inside sound-attenuating chambers located in an experimental room (ambient temperature 22±1 °C; illuminated by red light) outside of the housing vivarium. To begin a session, the catheter fittings on the animals' backs were connected to polyethylene tubing contained inside a protective spring suspended into the operant chamber from a liquid swivel attached to a balance arm. Each operant session started with the extension of two retractable levers into the chamber. Following each completion of the response requirement (response ratio), a white stimulus light (located above the reinforced lever) signaled delivery of the reinforcer and remained on during a 20-sec post-infusion timeout, during which responses were recorded but had no scheduled consequences. Drug infusions were delivered via syringe pump, located outside of the sounds-attenuating chamber. The training dose (0.15 mg/kg/infusion; ~0.1 ml/infusion) was selected from prior self-administration studies (Nguyen et al., 2019; Wade, Vendruscolo, Schlosburg, Hernandez & Koob, 2015). The session duration for the Short Access (ShA) group was 1 h and the Long Access (LgA) training sessions were 12h (or 11h for ICSS-trained rats) in duration. Self-administration sessions were scheduled during weekdays with drug abstinence on the weekend.

*Progressive-Ratio (PR) Dose-Response Testing:* Following acquisition rats were subjected to randomized dose-substitution conditions under a Progressive Ratio (PR) schedule of reinforcement wherein different per-infusion doses of oxycodone (0, 0.06, 0.15, 0.3 mg/kg/inf) were presented in a balanced order on sequential sessions lasting up to 3 h. For the PR task, the sequence of response ratios started with one response then progressed through ratios determined by the following equation (rounded to the nearest integer): response ratio=5e^(injection number x j) – 5 (Richardson & Roberts, 1996). The value of 'j' was 0.2 and was chosen so as to observe a 'breakpoint' within ~3 h. The last ratio completed before the end of the session (1 h after the last response up to a maximum 3 h session) was operationally defined as the breakpoint. The dose order was balanced by Latin Square design (i.e. the 4 total conditions were randomized).

**Intracranial Self-Stimulation (ICSS) Reward Procedure.** Rats (N=34) were anesthetized (1-3% isoflurane in oxygen) and positioned in a stereotaxic frame (Kopf Instruments, Tujunga, CA). The incisor bar was adjusted to 5 mm above the interaural line. Rats were prepared with unilateral electrodes aimed at the medial forebrain bundle (coordinates: AP -0.5mm, ML ±1.7mm, DVskull -9.5mm). Animals were trained in a procedure adapted from the discrete-trial current-threshold procedure (Kenny & Markou, 2006; Kornetsky & Esposito, 1979; Markou & Koob, 1992; Nguyen, Aarde, Cole, Vandewater, Grant & Taffe, 2016). Trials begin with a noncontingent stimulation (sinusoidal electrical stimuli of 250 ms duration and 60Hz), followed by a variable post-stimulation interval (7.5 s) during which delivery of a second stimulus was contingent upon responding with a ¼ turn of a wheel manipulandum. Each electrical stimulation (reinforcer) had a train duration of 500 ms during which 0.1 ms cathodal pulses were delivered at 50-100 Hz, with current-intensity thresholds within 50-200 µA. Current was varied in a series of steps (±5 µA per step, 3 trials per step). In each testing session, four alternating descending-ascending series were presented. The threshold for each series was defined as the midpoint between two consecutive current intensities that yielded ‘positive scores’ (animals responded for at least two of the three trials) and two consecutive current intensities that yielded ‘negative scores’ (animals did not respond for two or more of the three trials). The overall threshold of the session was defined as the mean of the thresholds for the four individual series. Each testing session was ~30 min in duration. Rats were trained once daily until stable reward threshold were established (<10% variation in thresholds for three consecutive days) between 7 and 10 days. Following ICSS training, rats were implanted with intravenous catheters (see above). After a 7 day recovery, the rats were tested in the ICSS experiments for 5 days without drug exposure to determine baseline reward thresholds (average of the 5 days). Starting the following week ICSS sessions were run ~1 h prior to the start of each oxycodone self-administration session.

**Experiment 1: Intravenous Self-Administration of Oxycodone Under Short and Long Access Conditions.** Rats were randomly assigned to 12 h LgA (N=12) and 1 h ShA (N=12) groups. Two rats in the LgA group that failed to average more than 24 infusions (defined by the highest responder in a Saline LgA group, see (Nguyen et al., 2017)) over the final 5 sessions of acquisition were excluded from the acquisition and PR self-administration data as non-escalators. Three rats in the LgA group failed patency testing during the acquisition interval and were excluded. Rats were returned to their home cages for an extended 30-day abstinence period and then returned to 12 h sessions to test for re-engagement of drug seeking (re-escalation) following detoxification.

**Experiment 2: ICSS Reward During Intravenous Self-Administration of Oxycodone.** For these studies, rats completed daily (M-F) self-administration sessions after ICSS sessions for three sequential weeks, with the self-administration session omitted on the Thursday of Week 3. Six rats were randomly assigned to ShA (1 h; N=6) and twelve rats to LgA (11 h; N=12) since the primary goal was to assess LgA IVSA. The 11 h LgA self-administration duration was selected to preserve the interval of discontinuation (12h) from behavior testing to match the IVSA-only cohort. The effect of non-contingent oxycodone (see below) was assessed during Weeks 4-5 and again in Week 9. Week 6 constituted an uninterrupted week of ICSS followed by standard IVSA sessions. Self-administration sessions were omitted during Week 7 to assess protracted discontinuation of drug access. In Week 8, rats received ICSS evaluation before and after one hour of IVSA on Monday and Friday (the remaining 10 h of self-administration were omitted for the LgA groups on those days). The LgA rats were switched to 4 h daily IVSA sessions for Week 9 and received no self-administration sessions in Week 10. One rat was excluded from the study due to opioid-induced self-injury during acquisition. Two rats were excluded due to technical failure of the catheter, one during acquisition and one at the start of Week 8.

*Non-contingent drug exposure:* During Weeks 11-12, rats received acute doses of methamphetamine (0.5 mg/kg, i.p.; Week 11) or oxycodone (0.25 mg/kg, s.c.; Week 12) 15 min prior to the ICSS session to determine if brain reward sensitivity to rewarding drug administration was restored after sustained discontinuation. Doses and routes of administration were selected based upon approximate conditions previously shown to significantly facilitate ICSS responding (Nguyen, Aarde, Cole, Vandewater, Grant & Taffe, 2016; Wiebelhaus, Walentiny & Beardsley, 2016).

**Experiment 3: Effect of KOR antagonism on ICSS Reward Threshold in Rats Trained to Self-administer Oxycodone Under Extended Access Conditions.** Separate groups of rats were trained in the ICSS procedure, as above, and administered norBNI (30 mg/kg, i.p.; N=8, LgA-norBNI) or saline vehicle (N=8, LgA-sal) 3 days prior to oxycodone self-administration training. One LgA-norBNI rat did not exhibit the threshold lowering effect of methamphetamine on ICSS and was therefore excluded. During Weeks 3 and 6, rats received a second ICSS evaluation after one hour of oxycodone self-administration on Friday (the remaining 10 h of self-administration were omitted for the LgA groups on those days). In Weeks 6-7, rats were tested under continuous daily 11 h oxycodone IVSA sessions for eight days and thereafter switched to daily saline IVSA for the next 4 sessions to determine the impact of drug abstinence, while leaving all other IVSA procedures intact. In Week 8, rats had access to oxycodone IVSA for only 1 h to assess the impact of ShA versus LgA within-group. In Weeks 9-10, rats underwent heroin (0.06 mg/kg/infusion) substitution experiments for 11 and 4 h to determine if effects generalized to a different opioid. During Week 9, rats received a second ICSS evaluation after one hour of heroin self-administration on Friday (the remaining 10 h of self-administration were omitted for the LgA groups on those days). In Weeks 11-12 rats received THC (5 mg/kg, i.p.) or vehicle after the ICSS session and 30 minutes prior to the start of IVSA for four sequential days. The order of THC vs vehicle treatment was counterbalanced across the two weeks. Analyses of these data included mixed-effects models in Prism (Graphpad Software, Inc, La Jolla, CA).

### Supplementary Results and Discussion

<u>Comparisons of initial oxycodone intake under long (11 h) or short (1 h) access conditions</u>

Analysis of the first 10, 30 and 60 minute intakes for Experiment 1 shows that by Session 15, the criterion of increased intake in the LgA group versus the ShA group, under standard training conditions, was met (**Figure S1**). In these analyses, the 30 minute data are inclusive of the 10 minute data, and the 60 minute data are inclusive of the 10 and 30 minute data. Some investigations describing escalated self-administration of heroin include the first hour comparison (Ahmed, Walker & Koob, 2000; Greenwell et al., 2009), whereas other papers show that there was no difference in first hour intake (Greenwell, Walker, Cottone, Zorrilla & Koob, 2009; Park, Schlosburg, Vendruscolo, Schulteis, Edwards & Koob, 2015; Schmeichel et al., 2015; Vendruscolo, Schlosburg, Misra, Chen, Greenwell & Koob, 2011). An additional set of studies fail to present the first hour intakes for the extended-access groups that were shown/tested (Edwards et al., 2012; McFalls, Imperio, Bixler, Freeman, Grigson & Vrana, 2016; McNamara, Dalley, Robbins, Everitt & Belin, 2010; Picetti, Caccavo, Ho & Kreek, 2012; Schlosburg et al., 2013; Wade, Vendruscolo, Schlosburg, Hernandez & Koob, 2015) which might lead to speculation that there was indeed no difference in the first hour under standard training conditions.

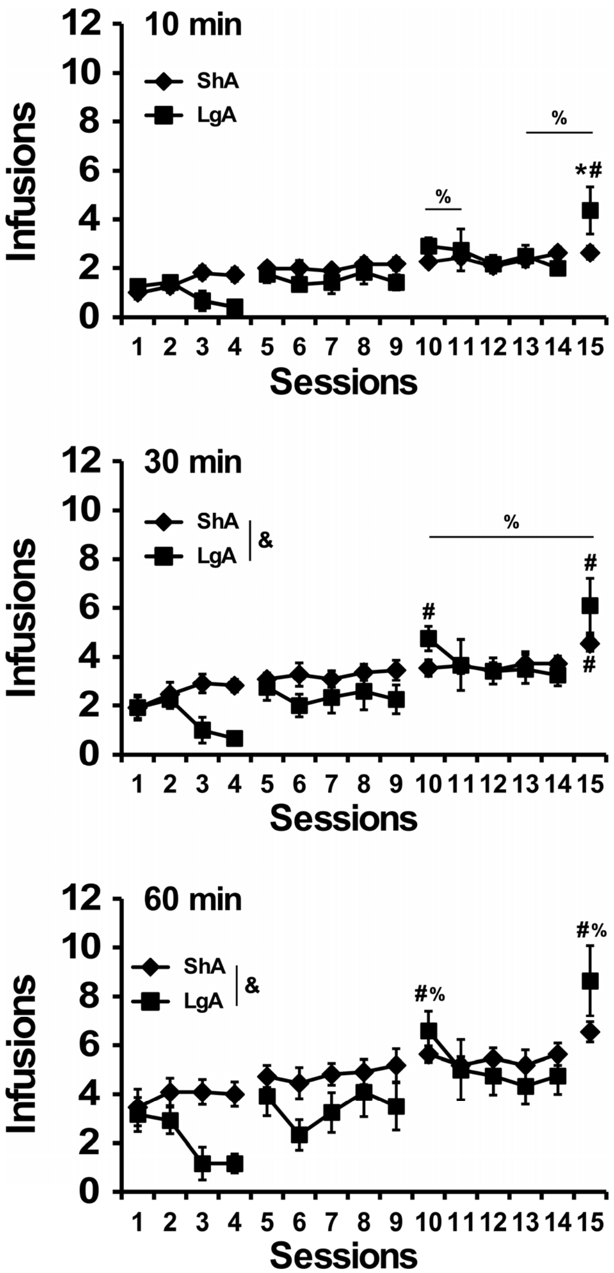

***Figure S1. Time bin analysis of oxycodone self-administration under LgA and ShA conditions.*** *Mean (±SEM) oxycodone infusions obtained by LgA (N=12) and ShA rats (N=11) during the initial 10, 30 or 60 minutes of a self-administration session. Significant difference in oxycodone infusions between LgA and ShA rats is indicated by *, from Session 1 within-group is indicated by #, and from Session 1 across group means is indicated by %, Difference in group means is indicated by &.*

Reward threshold elevations following non-contingent drug administration

The elevation in reward threshold that was observed later in the self-administration weeks could be reversed by a pre-session injection (0.0, 0.25, 0.5, 1.0 mg/mkg, i.p.) of oxycodone in a dose-dependent manner (**Figure S2**). Analysis confirmed a significant effect of Dose [F(4,36)=6.678; *p*<0.0001] and of the interaction of Dose X Withdrawn/IVSA factor [F{1,9)=2.425; *p*<0.005]. Post hoc analysis confirmed reward thresholds were significantly difference between LgA-withdrawn and LgA-IVSA following injections of saline vehicle and 0.25 mg/kg, s.c. oxycodone. ICSS thresholds were significantly higher following saline and 1.0 mg/kg, s.c. oxycodone in the LgA-IVSA group compared to baseline, and thresholds were higher following1.0 mg/kg s.c. oxycodone in the LgA-withdrawan group compared to all other dose conditions.

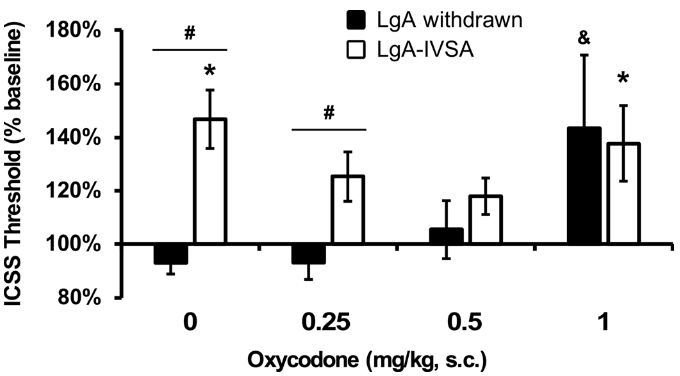

***Figure S 2. Non-contingent oxycodone normalizes elevated brain reward thresholds.***
*Mean (N=10; ±SEM) ICSS thresholds following systemic injection of oxycodone following 60 h discontinuation (LgA withdrawn) and 5 sessions oxycodone self-administration (LgA-IVSA) sessions. The data are expressed as percent of the individual baseline threshold. A significant difference from baseline is indicated with *, a difference between LgA-IVSA and LgA-withdrawn dose series by # and a difference from all other dose conditions by &.*

Following 2-3 week discontinuation of intravenous self-administration, additional studies analyzed reward threshold in 7 remaining rats (4 rats were excluded due to unstable ICSS performance) during Weeks 11-12 (**Figure S3**). Reward thresholds were significantly reduced 15 min after acute injection of methamphetamine (0.5 mg/kg, i.p.; 3 weeks discontinuation) or oxycodone (0.25 mg/kg. s.c.; 4 weeks discontinuation). Analysis confirmed a significant effect of Drug Treatment

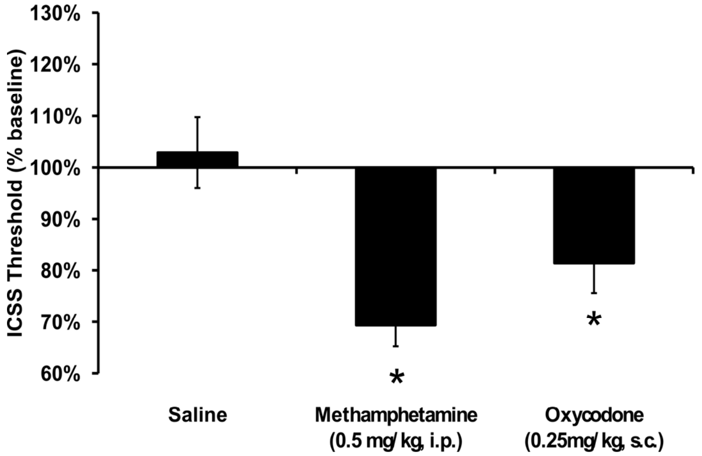

***Figure S3. Decrease in Brain Reward Thresholds from Methamphetamine or Oxycodone Injection following Extended Discontinuation.***
*Mean (N=7; ±SEM) ICSS thresholds following challenge with single doses of methamphetamine or oxycodone. The data are expressed as percent of the individual baseline threshold. A significant difference from the Saline is indicated with *.*

[F(1.257,7.541)=8.72; *p*<0.05] and the post hoc test confirmed reward thresholds were significantly lower following injections of both drugs compared to injection of saline.

Bi-directional investigation of brain reward thresholds and oxycodone self-administration behavior

The intracranial self-stimulation reward procedure failed to affect oxycodone self-administration (**Figure S4**). To determine if the ICSS procedure prior to the IVSA sessions impacted oxycodone intake, the acquisition data for the four LgA cohorts was compared using the days of access that were shared across all four groups. This meant that the fourth day of the third week was omitted (because it was omitted for the animals in Experiment 2), the fifth day of the third week was omitted (because it was only a one hour session for the Experiment 3 groups) and one day of the first week was omitted because Experiment 1 started on a four day week. This analysis did not confirm any effect of Group [F(3,29)=0.41; *p*=0.7442] or any interaction of Group with Session [F(36,348)=1.08; *p*=0.3466] on the number of infusions obtained. There was however a significant effect of Session [F(12,348)= 8.86; *p*<0.0001]. During Sessions 51-54 for the Experiment 3 groups, the ICSS session was omitted twice, and included twice, in counter balanced order to determine any effects on oxycodone self-administration. No significant effect of omitting the ICSS session was confirmed.

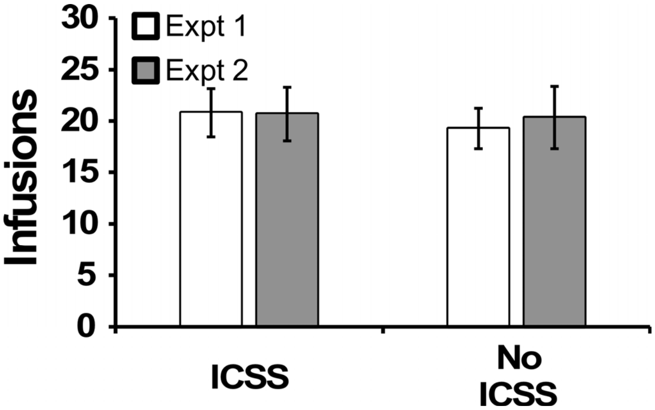

***Figure S4. Self-administration of oxycodone under 2 h access conditions with and without prior ICSS-training.***
*Mean (±SEM) oxycodone infusions obtained by LgA (N=8) and LgA-norBNI (N=7) rats under 2 h access conditions. The IVSA sessions were run after a normal ICSS session or without any prior ICSS that day, in a balanced order on two successive weekdays. The experiment was repeated twice across an interval of a week. A significant difference from the Session 1 is indicated with *.*

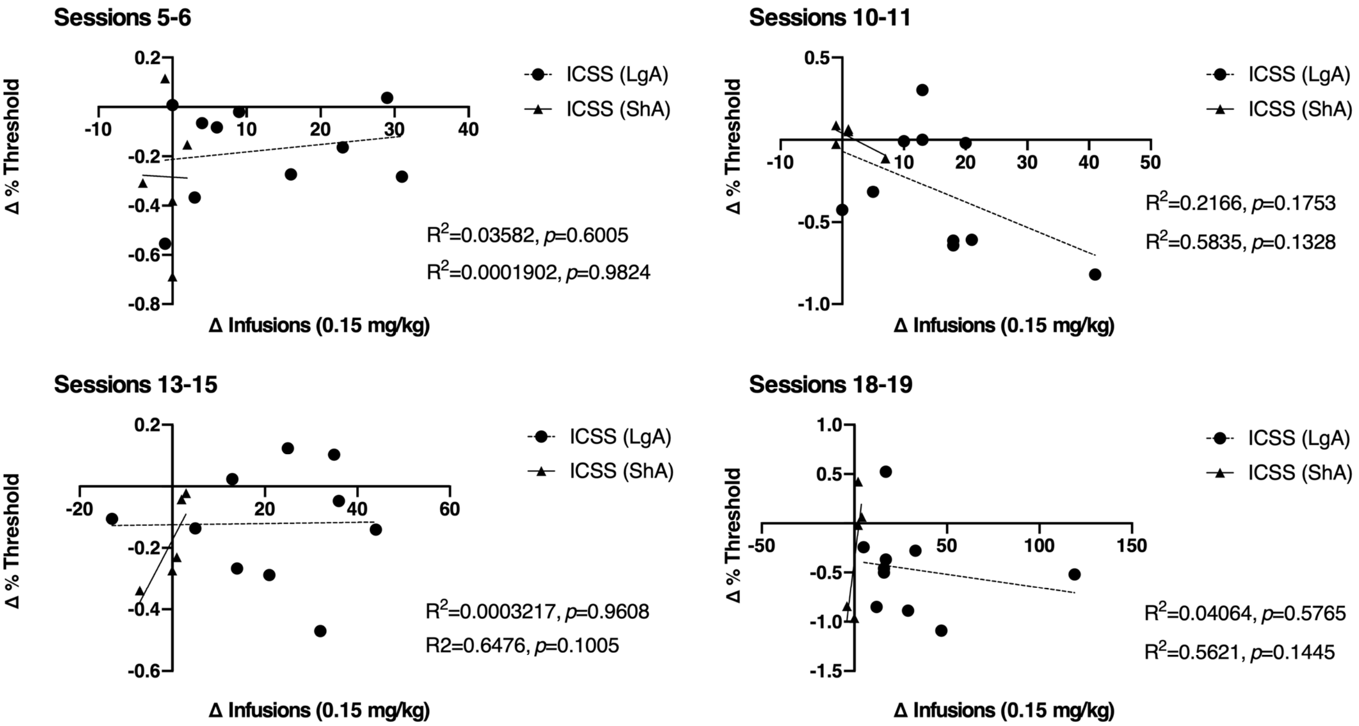

***Figure S5. Correlational Analysis of the relationship between oxycodone infusions and changes in ICSS brain reward threshold following weekend abstinence.***
The relationship between intravenous oxycodone infusions and ICSS brain reward thresholds show a lack of Pearson correlation ($R^2 < 0.6476$; $p > 0.05$). Reward thresholds were analyzed using the change between Pre- and Post-weekend (60 h) abstinence from self-administration (during Sessions 5-6. 10-11, 13-15, 18-19 respectively). Infusions (IVSA) were collected from rats trained under either long (LgA) or short (ShA) access conditions.

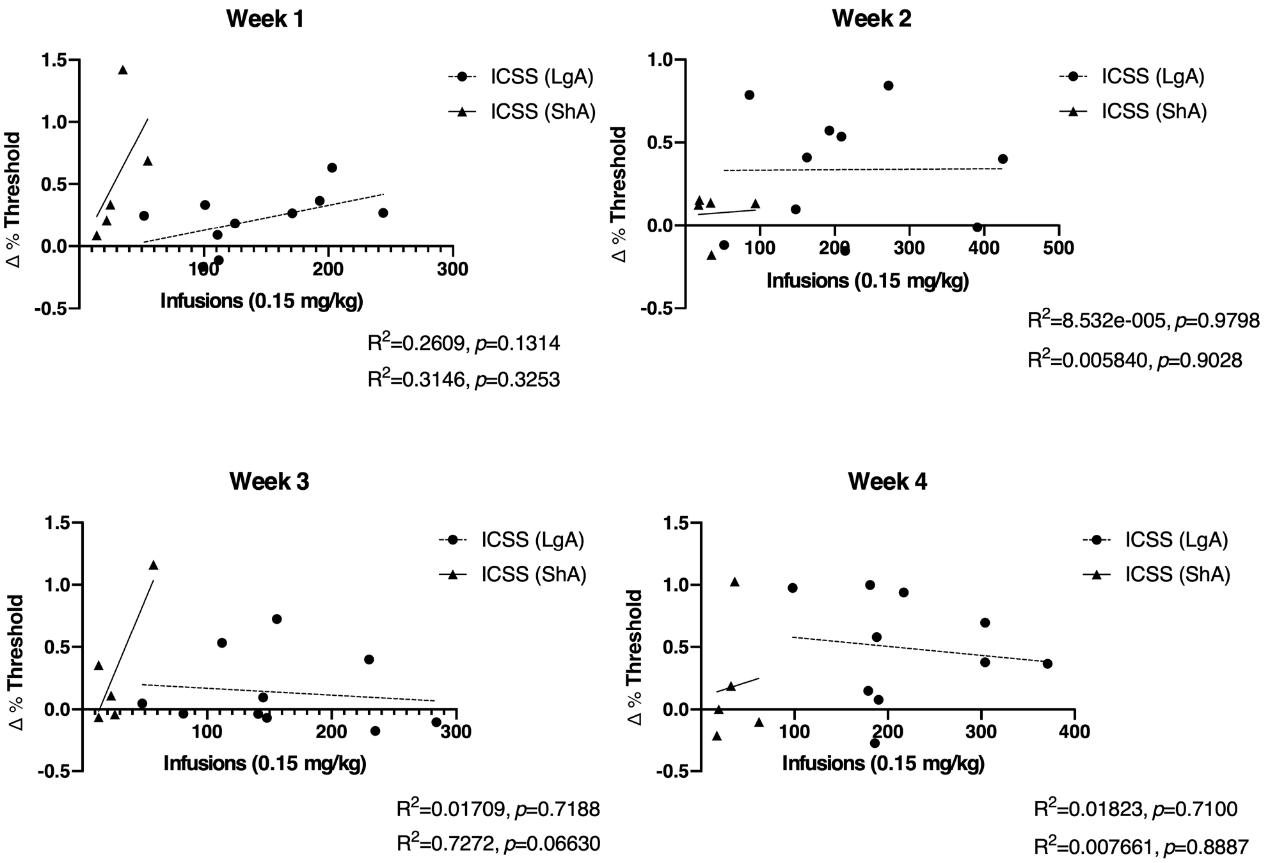

***Figure S6. Correlational Analysis of the relationship between cumulative oxycodone infusions and changes in ICSS brain reward threshold.***

The relationship between intravenous oxycodone infusions and ICSS brain reward thresholds show a lack of Pearson correlation ($R^2 < 0.7272$; $p > 0.05$). Weekly change in reward thresholds (Monday to end of the week) were analyzed using the cumulative oxycodone infusions (Sessions 1-5, 6-10, 11-13, 15-18 respectively). Infusions (IVSA) were collected from rats trained under either long (LgA) or short (ShA) access conditions.

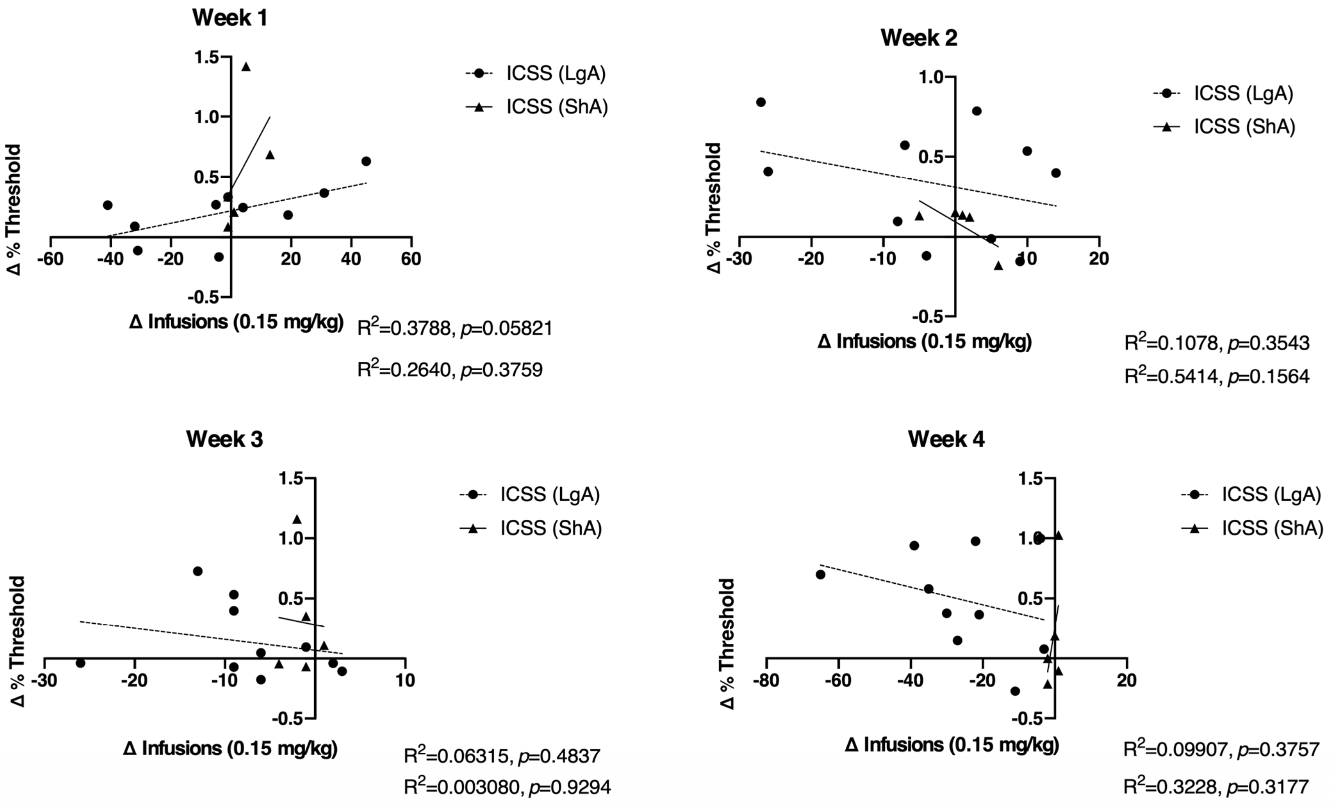

***Figure S7. Correlational Analysis of the relationship between within week changes in oxycodone infusions and changes in ICSS brain reward.***

The relationship between intravenous oxycodone infusions and ICSS brain reward thresholds show a lack of Pearson correlation ($R^2 < 0.5414$; $p > 0.05$). Weekly change in reward thresholds (Monday to end of the week) were analyzed using the change in oxycodone infusions (differences between Sessions 1 and 5, 6 and 10, 11 and 13, 15 and 18 respectively). Infusions (IVSA) were collected from rats trained under either long (LgA) or short (ShA) access conditions.

**Expanded Results Narrative with Analysis Details**

**Escalation of oxycodone self-administration under extended access conditions**

In Experiment 1, the mean number of intravenous oxycodone infusions obtained by rats trained under LgA (N=8) conditions was significantly higher than infusions obtained by ShA (N=12) rats (**Figure 1A**). The ANOVA confirmed significant main effects of Session [F(14,252)=24.3; $p<0.0001$], of Access Duration [F(1,18)=97.07; $p<0.0001$], and of the interaction of factors [F(14,252)=21.16; $p<0.0001$]. Post hoc analysis confirmed that more infusions were obtained by LgA rats during sessions 5 and 7-15 compared to session 1, LgA rats received significantly more infusions compared to ShA rats during sessions 5-15 and that drug intake did not change across sessions for the ShA group. Analysis of the first 10, 30 and 60 minutes of self-administration confirmed escalation in the LgA group but not in the ShA group (see **Supplementary Figure S1**). During the final session of acquisition, LgA rats exhibited 76.9% drug-associated lever responding. The ShA group exhibited >80% drug-associated lever responding for 9 of the final 11 days of acquisition. Analysis of the sessions before and after the 60 h weekend abstinence periods showed that LgA rats significantly increased drug intake following this extended drug deprivation, whereas ShA showed no significant change (**Figure 1B**). The LgA and ShA rats also exhibited group differences during PR dose substitution (**Figure 1C**) and the ANOVA confirmed significant main effects of Access [F(1,72)=10.77; $p<0.01$] and of Dose [F(3,72)=3.570; $p<0.05$]. Neither LgA (N=7) nor ShA (N=12) rats exhibited any significant change in oxycodone infusions (0.15 mg/kg/inf) obtained under a FR1 response contingency (**Figure 1D**) before versus after the 30-day abstinence interval. One rat (LgA) that completed the acquisition interval was not included in the post- abstinence assessment due to illness. The ANOVA confirmed a significant main effect of Access [F(1,17)=96.26; $p<0.0001$] and the post hoc analysis confirmed that significantly more infusions were obtained by LgA rats on the day after a weekend break.

**Brain stimulation reward during escalation of oxycodone self-administration under extended access conditions**

The LgA and ShA rats in the Experiment 2 (ICSS-trained) cohort also exhibited group differences in oxycodone self-administration (**Figure 2A**). The ANOVA confirmed significant effects of Session [$F(27,351)=3.99$; $p<0.0001$], of Group [$F(1,13)=19.89$; $p<0.001$] and of the interaction of factors [$F(27,351)=3.15$; $p<0.0001$] on the number of infusions obtained. The post hoc test confirmed that more infusions were obtained by the LgA group in sessions 14, 15, 19-24, 26-28 compared with the first session. In addition, significant group differences were confirmed across weekend breaks (Sessions 11-17 and 19-28) (**Figure 2B**). For LgA rats (N=10) the analysis of the weekend effects confirmed significant effects of Week [$F(4,36)=10.23$; $p<0.0001$] and of the Pre-/Post-weekend factor [$F(1,9)=26.56$; $p<0.001$] on oxycodone infusions. Post hoc test confirmed significant increases for each Monday session over the prior end of week session from the 11$^{th}$ session to the 24$^{th}$. Whereas for ShA rats (N=5), mean infusions were not significantly different between Pre-weekend and Monday sessions (**Figure 2C**).

ICSS thresholds in the LgA group in Experiment 2 increased across successive days in each week of self-administration but returned nearly to baseline across the weekend breaks (**Figure 3A**). The ANOVA confirmed a significant effect of Day [$F(4,36)=9.56$; $p<0.0001$] and of Week [$F(4,36)=5.91$; $p<0.001$] and of the interaction of factors [$F(16,144)=2.00$; $p<0.05$] on ICSS thresholds. The post hoc test further confirmed significant increases within each week of self-administration, no change within the abstinence week and differences between days of the self-administration weeks compared with the corresponding weekday of the abstinence week. An attenuation of the ICSS threshold elevation was observed on the 5th day of the third week, following omission of the IVSA session the previous day (see **Figure 2A**).

In Experiment 3, groups of rats (LgA-norBNI) that were injected with norBNI (30 mg/kg, i.p.), a kappa opioid receptor (KOR) antagonist, or saline (LgA-sal), prior to initiating self-administration training were used to determine the effect of KOR function on oxycodone-induced effects on ICSS reward thresholds (**Figure 3B**). Compared to LgA-sal control rats, LgA-norBNI rats had lower ICSS thresholds across successive days (Monday through Friday) during 4 weeks of self-administration. The analysis confirmed a significant effect of Session [$F(19,247)=6.949$; $p<0.0001$] and of Group [$F(1,13)=5.196$; $p<0.05$] but not the interaction of factors. Post hoc analysis confirmed ICSS thresholds were significantly increased in LgA-sal rats (Days 5, 9 and 18-20) and LgA-norBNI (Day 14) compared to the first Day of each Week. Interestingly, mean oxycodone infusions did not differ between LgA-sal control rats or LgA-norBNI rats (**Figure 3C**). Session 15 was excluded from this analysis, as the self-administration session was only 60 minutes followed by an ICSS session. The ANOVA confirmed a significant effect of Session [$F(18,234)=4.781$; $p<0.0001$] but not of Group or the interaction of factors. Post hoc analysis confirmed infusions were significantly increased in LgA-sal rats (Day 8,11,12, 14, 16, 19 and 20) compared to the first day of training; however, infusions were not significantly increased relative to Day 1 in LgA-norBNI rats.

We further confirmed in Experiment 3 that the within-week elevations of brain reward thresholds depended on access duration and generalized across oxycodone and heroin (**Figure 4A**). ICSS thresholds were elevated following seven sequential IVSA sessions (Days 16-23) in both LgA-sal and LgA-norBNI rats but returned to the baseline following four consecutive 11 h sessions of saline self-administration (Days 24-27). Analysis confirmed a significant effect of Session [F(7,91)=11.07; $p<0.0001$]. Subsequent oxycodone IVSA in 1 h access sessions in Week 6 (Days 28-31) failed to significantly increase reward thresholds. Heroin (0.06 mg/kg/infusion) IVSA under 11 h access duration (Days 32-41) significantly elevated reward thresholds in both groups [F(4,52)=22.6; $p<0.0001$] in Week 7 and then in 4 h access sessions in Week 8 [F(4,52)=2.587; $p<0.05$]. The Dunnett post hoc test, collapsed across groups, confirmed significant changes relative to pre-saline (Day 23) and the first day of each heroin week, respectively. Group comparisons of daily ICSS thresholds showed consistency across the LgA (Experiment 2) and LgA-saline (Experiment 3) groups, as they exhibited very similar elevations of reward thresholds across the initial three weeks (**Figure 4B**). More interestingly, the pattern of ICSS threshold increases in the LgA-norBNI (Experiment 3) group was very similar to the pattern of modest increases observed in the ShA (Experiment 2) group. Analysis comparing the ICSS threshold changes within each of the four groups across the first 14 sessions confirmed a main effect of group (F (3, 28) = 3.07; P<0.05] and of Session [F (13, 364) = 10.52; P<0.0001]. The Dunnett post hoc test of the marginal means did not confirm any significant difference between LgA and ShA ($p$=0.066) or LgA-norBNI ($p$=0.074) groups.

**Brain stimulation reward procedure does not affect oxycodone self-administration**

To determine if the ICSS procedure prior to the IVSA sessions impacted oxycodone intake, the acquisition data for the four LgA cohorts was compared using the days of access that were shared across all four groups. This meant that the fourth day of the third week was omitted (because it was omitted for the animals in Experiment 2), the fifth day of the third week was omitted (because it was only a one hour session for the Experiment 3 groups) and one day of the first week was omitted because Experiment 1 started on a four day week. This analysis did not confirm any effect of Group [$F(3,29)=0.41$; $p=0.7442$] or any interaction of Group with Session [$F(36,348)=1.08$; $p=0.3466$] on the number of infusions obtained. There was however a significant effect of Session [$F(12,348)= 8.86$; $p<0.0001$]. During Sessions 51-54 for the Experiment 3 groups, the ICSS session was omitted twice, and included twice, in counter balanced order to determine any effects on oxycodone self-administration. No significant effect of omitting the ICSS session was confirmed (see **Supplementary Figure S4**).

**Normalization of elevated ICSS thresholds by a one hour self-administration session**

Reward thresholds were assessed before and after a 1 h IVSA session in LgA groups in Experiments 2 (**Figure 5A**) and 3 (**Figure 5B-D**). In the Experiment 2 group, the analysis of the ICSS thresholds before and after a 1 h oxycodone IVSA session on Monday and Friday confirmed a significant effect of the Day of the week [$F(1,8)=39.29$; $p<0.001$] and a significant effect of the Pre/Post Session factor [$F(1,8)=5.334$; $p<0.05$]. The post hoc test confirmed that pre-IVSA reward thresholds were significantly higher on Friday compared with Monday and that a significant post-IVSA reduction in threshold was observed on Friday (**Figure 5A**). Analysis of ICSS thresholds after a 1 h oxycodone IVSA session in Experiment 3 (**Figure 5B**), confirmed there was a significant effect of Pre/Post Session [$F(1,13)=18.22$; $p<0.001$]. Post hoc analysis further confirmed a significant reduction in ICSS threshold in both LgA and LgA-norBNI groups. In contrast, there was no significant effect of the Pre/Post Session factor during daily 1 h oxycodone IVSA (**Figure 5C**). Finally, there was a significant effect of Pre/Post Session [$F(1,13)=23.16$; $p<0.001$] following 1 h IVSA of heroin during the daily 11 h heroin IVSA training (**Figure 5D**). Post hoc analysis confirmed a significant reduction in ICSS threshold in both LgA and LgA-norBNI groups.

**Combined reinforcer efficacy versus overall oxycodone intake**

In this study rats were injected daily with THC (5 mg/kg, i.p.) or vehicle prior to 6 h IVSA sessions for four sequential days; pretreatments were counterbalanced across two weeks within the total sample (i.e., LgA-sal and LgA-norBNI combined). Our previous study showed that THC decreases oxycodone self-administration (Nguyen et al., 2019) by enhancing the rewarding efficacy of a given unit dose. Data from the current experiment confirmed that this reduction in oxycodone self-administration behavior caused by THC (**Figure 6A**) did not change the within-week increase in ICSS thresholds (**Figure 6B**). Analysis of the self-administration data confirmed a significant effect of Treatment [F(1,15)=20; $p$<0.0005] but not of Session or the interaction of factors. Post hoc analysis confirmed that infusions were significantly different between rats injected with vehicle (Veh) or THC (5 mg/kg, i.p.) during days 2 and 3.

Analysis of ICSS data confirmed that reward thresholds significantly increased in the rats when treated with either vehicle or THC (**Figure 6B**). The ANOVA confirmed a significant main effect of Session [F3,84)=18.18; $p$<0.0001] but not of Treatment or of the interaction of factors. Interestingly, these data complement the original effect of norBNI in this group in which a difference in the ICSS pattern was observed without any difference in oxycodone self-administration, and further illustrate the partial independence of these indices. The THC effect is likely due to an increase in the reinforcing efficacy of a unit dose of oxycodone, as in our prior study. As the bin analysis of the self-administration pattern shows (**Figure 6C**), the THC condition blunts, but does not eliminate, intake during the loading phase and the first several hours. The rats do exhibit a loading phase and continue to respond for drug infusions in a regular pattern. Thus, THC is not merely suppressing behavior or, likely, the drug-seeking.

**References**

Edwards S, Vendruscolo LF, Schlosburg JE, Misra KK, Wee S, Park PE*, et al.* (2012). Development of mechanical hypersensitivity in rats during heroin and ethanol dependence: alleviation by CRF(1) receptor antagonism. Neuropharmacology 62**:** 1142-1151.

Greenwell TN, Walker BM, Cottone P, Zorrilla EP, & Koob GF (2009). The alpha1 adrenergic receptor antagonist prazosin reduces heroin self-administration in rats with extended access to heroin administration. Pharmacol Biochem Behav 91**:** 295-302.

Kenny PJ, & Markou A (2006). Nicotine self-administration acutely activates brain reward systems and induces a long-lasting increase in reward sensitivity. Neuropsychopharmacology 31**:** 1203-1211.

Kornetsky C, & Esposito RU (1979). Euphorigenic drugs: effects on the reward pathways of the brain. Fed Proc 38**:** 2473-2476.

Markou A, & Koob GF (1992). Construct validity of a self-stimulation threshold paradigm: effects of reward and performance manipulations. Physiology & Behavior 51**:** 111-119.

McFalls AJ, Imperio CG, Bixler G, Freeman WM, Grigson PS, & Vrana KE (2016). Reward devaluation and heroin escalation is associated with differential expression of CRF signaling genes. Brain Res Bull 123**:** 81-93.

McNamara R, Dalley JW, Robbins TW, Everitt BJ, & Belin D (2010). Trait-like impulsivity does not predict escalation of heroin self-administration in the rat. Psychopharmacology (Berl) 212**:** 453-464.

Nguyen JD, Aarde SM, Cole M, Vandewater SA, Grant Y, & Taffe MA (2016). Locomotor Stimulant and Rewarding Effects of Inhaling Methamphetamine, MDPV, and Mephedrone via Electronic Cigarette-Type Technology. Neuropsychopharmacology 41**:** 2759-2771.

Nguyen JD, Grant Y, Creehan KM, Hwang CS, Vandewater SA, Janda KD*, et al.* (2019). Delta(9)-tetrahydrocannabinol attenuates oxycodone self-administration under extended access conditions. Neuropharmacology 151**:** 127-135.

Nguyen JD, Grant Y, Creehan KM, Vandewater SA, & Taffe MA (2017). Escalation of intravenous self-administration of methylone and mephedrone under extended access conditions. Addiction biology 22**:** 1160-1168.

Nguyen JD, Kirson D, Steinman MQ, Patel R, Khom S, Varodayan FP*, et al.* (2017). Withdrawal-induced escalated oxycodone self-administration is mediated by kappa opioid receptor function. bioRxiv.

Park PE, Schlosburg JE, Vendruscolo LF, Schulteis G, Edwards S, & Koob GF (2015). Chronic CRF1 receptor blockade reduces heroin intake escalation and dependence-induced hyperalgesia. Addict Biol 20**:** 275-284.

Picetti R, Caccavo JA, Ho A, & Kreek MJ (2012). Dose escalation and dose preference in extended-access heroin self-administration in Lewis and Fischer rats. Psychopharmacology (Berl) 220**:** 163-172.

Richardson NR, & Roberts DC (1996). Progressive ratio schedules in drug self-administration studies in rats: a method to evaluate reinforcing efficacy. Journal of Neuroscience Methods 66**:** 1-11.

Schlosburg JE, Whitfield TW, Jr., Park PE, Crawford EF, George O, Vendruscolo LF*, et al.* (2013). Long-term antagonism of kappa opioid receptors prevents escalation of and increased motivation for heroin intake. J Neurosci 33**:** 19384-19392.

Schmeichel BE, Barbier E, Misra KK, Contet C, Schlosburg JE, Grigoriadis D*, et al.* (2015). Hypocretin receptor 2 antagonism dose-dependently reduces escalated heroin self-administration in rats. Neuropsychopharmacology 40**:** 1123-1129.

Vendruscolo LF, Schlosburg JE, Misra KK, Chen SA, Greenwell TN, & Koob GF (2011). Escalation patterns of varying periods of heroin access. Pharmacol Biochem Behav 98**:** 570-574.

Wade CL, Vendruscolo LF, Schlosburg JE, Hernandez DO, & Koob GF (2015). Compulsive-like responding for opioid analgesics in rats with extended access. Neuropsychopharmacology 40**:** 421-428.

Wiebelhaus JM, Walentiny DM, & Beardsley PM (2016). Effects of Acute and Repeated Administration of Oxycodone and Naloxone-Precipitated Withdrawal on Intracranial Self-Stimulation in Rats. J Pharmacol Exp Ther 356**:** 43-52.
